# The STRIPAK complex is required for radial sorting and laminin receptor expression in Schwann cells

**DOI:** 10.1101/2024.10.30.620661

**Authors:** Michael R. Weaver, Dominika Shkoruta, Marta Pellegatta, Caterina Berti, Marilena Palmisano, Scott Ferguson, Edward Hurley, Julianne French, Shreya Patel, Sophie Belin, Matthias Selbach, Florian Ernst Paul, Fraser Sim, Yannick Poitelon, M. Laura Feltri

## Abstract

During peripheral nervous system development, Schwann cells undergo Rac1-dependent cytoskeletal reorganization as they insert cytoplasmic extensions into axon bundles to radially sort, ensheath, and myelinate individual axons. However, our understanding of the direct effectors targeted by Rac1 is limited. Here, we demonstrate that striatin-3 and MOB4 are novel Rac1 interactors. We show that, similar to Rac1-null Schwann cells, Schwann cell specific ablation of striatin-3 causes defects in lamellipodia formation. In addition, conditional Schwann cell knockout of multiple striatin proteins presents a severe delay in radial sorting. Finally, we demonstrate here that deletion of Rac1 or striatin-1/3 in Schwann cells causes defects in Hippo pathway regulation, phosphorylation of the Hippo pathway effectors YAP and TAZ, and expression of genes co-regulated by YAP and TAZ, such as extracellular matrix receptors. In summary, our results indicate that striatin-3 is a novel Rac1 interactor, show that striatin proteins are required for peripheral nervous system development, and reveal a role for Rac1 in regulation of the Hippo pathway in Schwann cells.

## INTRODUCTION

Radial sorting of axons by Schwann cells (SCs) in the peripheral nervous system (PNS) is a prerequisite for their proper myelination. Failure of radial sorting results in profound developmental defects. Surprisingly, most of the molecules that have been implicated in the radial sorting process are not at the apical, axoglial (adaxonal) interface, but on the basal (abaxonal) side of SCs. Indeed, the laminin-rich extracellular matrix (ECM), ECM receptors, and intracellular signaling triggered by ECM receptor-ligand engagement at the abaxonal interface are essential for SCs to radially sort axons (*Berti, et al., 2011; Court, et al., 2009; Feltri, et al., 2002; Feltri, et al., 2016; Nodari, et al., 2008; Nodari, et al., 2007; Occhi, et al., 2005; Pellegatta, et al., 2013; Previtali, et al., 2003a; Previtali, et al., 2003b; Saito, et al., 2003; Yu, et al., 2005*).

Signaling from laminin receptors also interacts with other signaling cascades that are essential for SC development. Notably, both remodeling of the actin cytoskeleton and cell polarity are regulated by Rho GTPase family members, and several Rho GTPases, including Rac1 and CDC42, are required for radial sorting (*Benninger, et al., 2007; Guo, et al., 2012; Nodari, et al., 2007*). Rac1 and CDC42 have been shown to regulate the activation of PKA through NF2. Moreover, expression of dominant-negative NF2 partially rescues radial sorting defects in CDC42 mutant mice, but not in Rac1 mutant mice (*Guo, et al., 2012; Guo, et al., 2013*). This suggests that the radial sorting defects observed in Rac1 mutant mice are mediated by additional mechanisms, and our understanding of the direct effectors targeted by Rac1 during radial sorting remains limited.

We previously showed that the Hippo pathway, a signaling cascade that negatively regulates the activity and nuclear translocation of the transcriptional co-activators YAP and TAZ, is also implicated in SC radial sorting (*Feltri, et al., 2021; Poitelon, et al., 2016*). YAP and TAZ notably regulate the expression of several laminin receptors and transcription factors essential for myelination (*Belin, et al., 2019; Deng, et al., 2017; Grove, et al., 2017; Hong, et al., 2024; Poitelon, et al., 2016*). It is well characterized that signals from the Hippo pathway, including from the kinases MST1 and MST2, lead to YAP and TAZ inhibition by phosphorylation and cytosolic sequestration. However, it is unknown what additional mechanisms may regulate YAP and TAZ in SCs.

Here, we sought to identify the downstream effectors of Rac1. We demonstrate that striatin-3 (STRN3), a member of striatin-interacting phosphatase and kinase (STRIPAK) complexes, directly interacts with Rac1. We found that loss of both striatin-1 (STRN1) and STRN3 in SCs leads to a severe delay in radial sorting. Finally, we demonstrate that loss of Rac1 or STRN1 and STRN3 in SCs leads to dysregulation of the Hippo pathway and altered expression of laminin receptors, thus revealing a novel role for Rac1 in SCs.

## RESULTS

### STRN3 and MOB4 both directly interact with Rac1

To screen for candidate Rac1 interactors, we performed a pulldown assay for active versus inactive forms of Rac1 from mouse peripheral nerve lysate at post-natal day (P)5, when both radial sorting and myelination are ongoing (*Feltri, et al., 2016*). Both STRN3 and MOB4, members of the STRIPAK complex, were among the proteins found to be enriched in the active, Rac1-GTP fraction (Fig. 1A-B). We confirmed that this interaction occurred specifically in SCs using both co-immunoprecipitation (co-IP) (Fig. 1C) and proximity ligation assay (PLA) (Supplementary Fig. 1).

**Figure 1:**
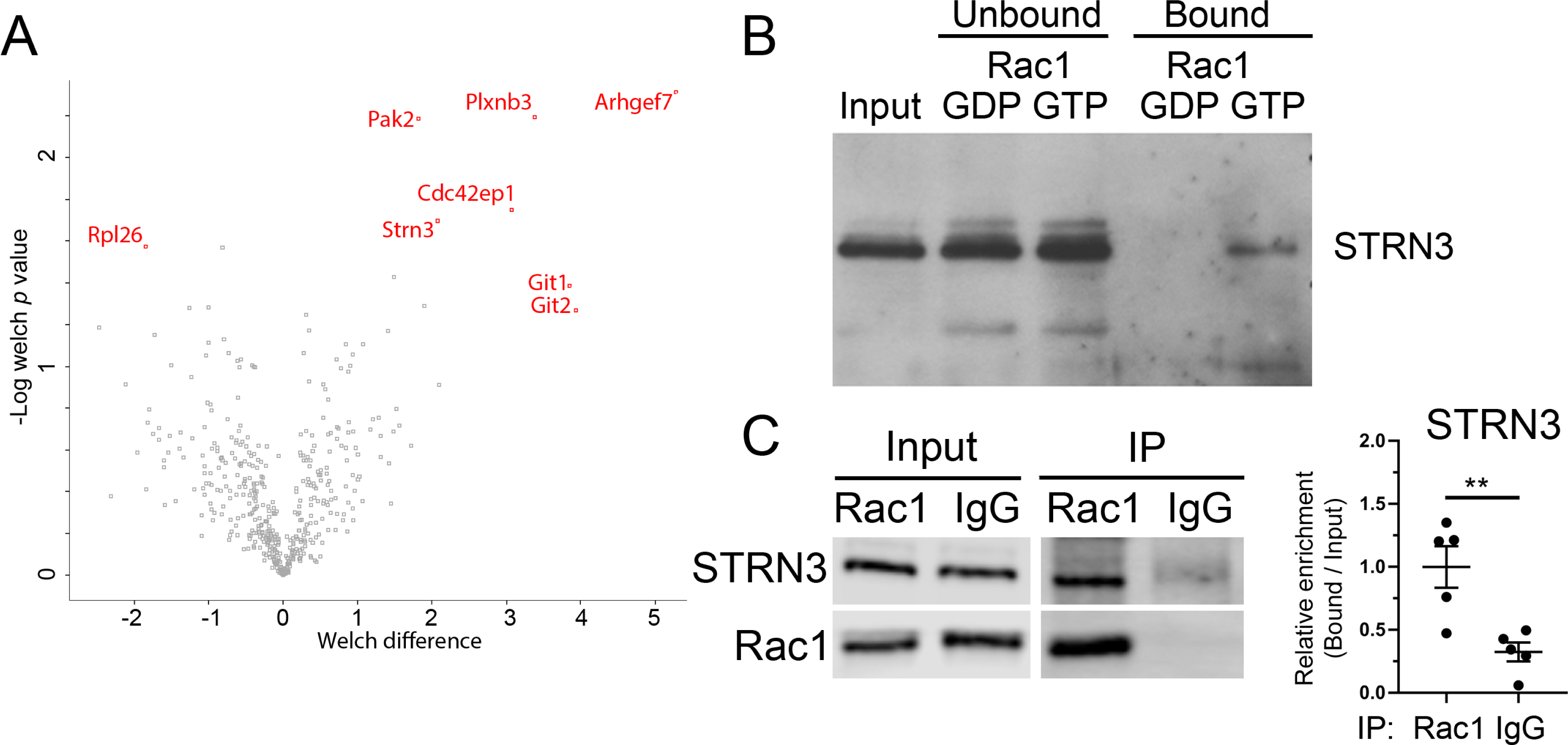
STRN3 interacts directly with Rac1. (**A**-**B**) GST pull-down from P5 wild-type (WT) mouse sciatic nerve lysate, comparing GST-Rac1-GDP vs GST-Rac1-GTP. (**A**) Volcano plot showing the Welch difference (log2 fold change) of Rac1 interactors. Proteins in red, such as STRN3 and several well-established Rac1 interactors, are significantly (FDR = 0.05) enriched in the active, Rac1-GTP fraction compared to the inactive, Rac1-GDP fraction. MOB4 was found to be enriched in the Rac1-GTP fraction in a separate biological replicate (not shown). (**B**) GST-Rac1-GDP/GTP pulldown of P5 WT sciatic nerve lysate was repeated and analyzed by western blot (WB). STRN3 was observed to elute with the active, Rac1-GTP fraction, but not with the inactive, Rac1-GDP fraction. (**C**) Immunoprecipitation of Rac1 from SC lysates. STRN3 was found to interact with Rac1 in SCs. *N* = 5 individual co-IPs per targeting antibody. Unpaired two-tailed *t*-test (*t* = 3.730, df = 8, *p* = 0.0058). Error bars indicate S.E.M. ** *p* < 0.01.

Although the Rac1 pulldown assay, co-IP, and PLA experiments strongly suggest that Rac1 associates with STRN3 in tissues and cells, they do not differentiate whether this interaction is direct or indirect. Therefore, we asked if STRN3 or MOB4 bind directly and specifically to Rac1, and in a Rac1-GDP/GTP charge-dependent manner. We used a series of dot blot assays with isolated proteins, which were all independently validated for purity and specificity (Supplementary Fig. 2-4). We found that both STRN3 and MOB4 bind directly to Rac1 (Supplementary Fig. 5). However, at low concentrations, and while isolated from the intracellular milieu, STRN3 exhibits stronger binding to inactive Rac1.

### STRN3 is required for SC elongation and process extension

We and others have previously demonstrated that Rac1 inhibition or ablation causes SCs to form fewer lamellipodia, a cytoplasmic extrusion that is critical for Rac1-mediated radial sorting and myelination (*Benninger, et al., 2007; Guo, et al., 2012; Nodari, et al., 2007*). Thus, we asked if loss of STRN3 in SCs may also influence lamellipodia formation and the radial sorting of axons. We generated mice lacking STRN3 in SCs (Strn3SCKO) using animals expressing Cre recombinase under control of the SC-specific *Mpz* promoter, which is expressed starting on embryonic day (E)13.5 (*Feltri, et al., 1999*). We show that SCs isolated from Strn3SCKO mice had little to no STRN3 protein expressed when compared to control SCs (Fig. 2A-B). Ablation of *Strn3* in SCs did not affect the purity of SC isolation (Fig. 2C-D) or the adherence of SCs to the substrate in vitro (Fig. 2E). However, similar to Rac1 inhibition or ablation in SCs, the *Strn3* null SCs had significantly fewer radial and total lamellipodia per cell (Fig. 2C, F) (*Nodari, et al., 2007*). We also found that *Strn3* null SCs had a significant defect in cell elongation (Fig. 2C, G-H). These data suggest that, like Rac1, STRN3 is a critical regulator of SC cytoskeletal dynamics, specifically, cell elongation, and lamellipodia formation.

**Figure 2:**
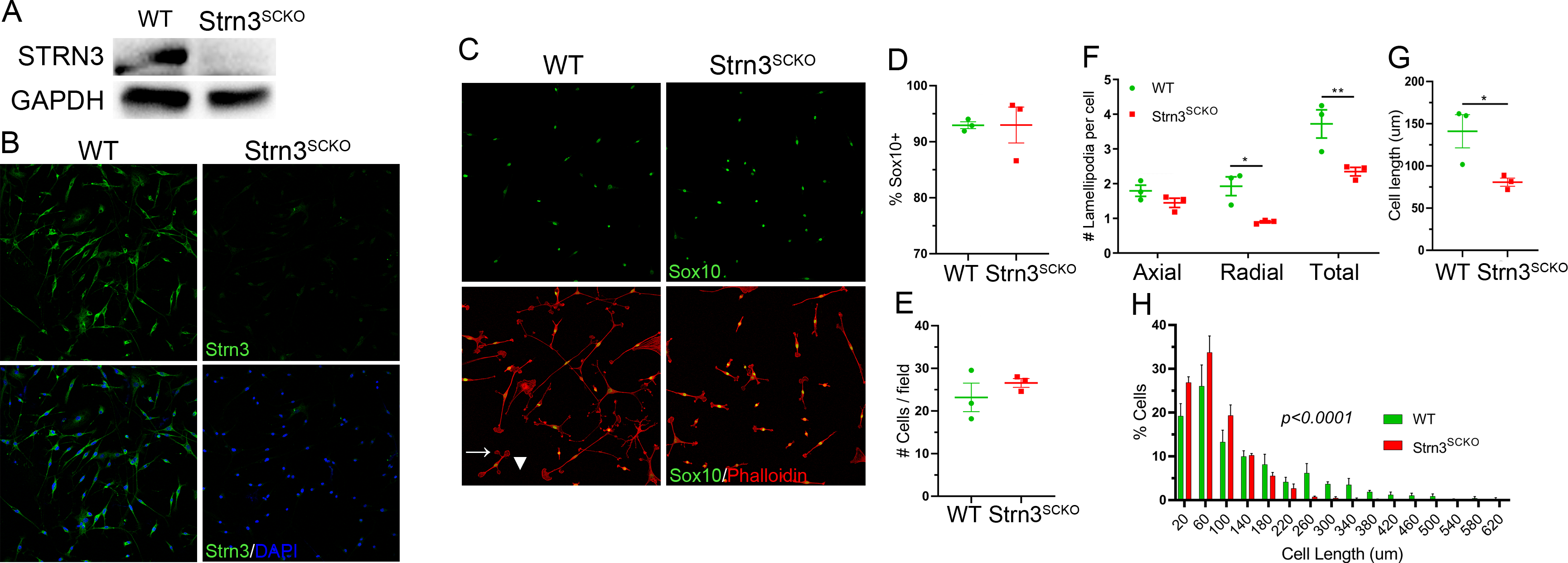
STRN3 is required for SC elongation and lamellipodia formation. SCs were isolated from WT and Strn3^SCKO^ littermates at P45-60. (**A**-**B**) SCs isolated from Strn3^SCKO^ mice have absent Strn3 protein expression by western blot (**A**) and immunohistochemistry (**B**). Similar results were obtained with a second anti-STRN3 antibody (data not shown). STRN3 was stained in green and nuclei were labeled with DAPI (blue). Scale bar = 20 µm. (**C**) Representative immunohistochemistry of mouse SCs, with nuclei labeled with Sox10 (green) and F-actin labeled with phalloidin (red). (**D**) Loss of STRN3 in SCs does not alter the purity of SC isolation. Unpaired two-tailed *t*-test (*t* = 0.01239, df = 4, *p* = 0.9907). (**E**) Deletion of *Strn3* in SCs does not affect the density of SCs adhering to the substrate in vitro. Unpaired two-tailed *t*-test (*t* = 0.9691, df = 4, *p* = 0.3874). (**F**) SCs isolated from Strn3^SCKO^ mice form reduced numbers of radial and total, but not axial, lamellipodia. Axial lamellipodia (white arrowhead) are defined as extending from within a 20° angle of the long axis of the cell, whereas radial lamellipodia (white arrow) protrude from outside of this range. Two-way multiple comparisons ANOVA with Bonferroni post hoc test. *F* (1,6) genotype = 45.63, *p* = 0.0005; *F* (2,6) lamellipodia type = 21.75, *p* = 0.0018; *p*_axial_ = 0.5662, *p*_radial_ = 0.0142, *p*_axial+radial_ = 0.0033. (**G**-**H**) SCs ablated for *Strn3* have reduced cell elongation. (**G**) Unpaired two-tailed *t*-test (*t* = 2.978, df = 4, *p* = 0.0408). (**H**) Lognormal nonlinear regression, extra sum-of-squares F test; *F* (3, 90) = 14.47. *N* = 3 individual mouse SC isolations per genotype. Error bars indicate S.E.M. * *p* < 0.05, ** *p* < 0.01.

### Striatin family protein expression is developmentally regulated

All three striatin proteins share a similar domain structure and a high degree of homology. Additionally, the main known function of each member of the striatin protein family is to act as the center scaffold of STRIPAK complexes (*Castets, et al., 1996; Castets, et al., 2000; Goudreault, et al., 2009*). Accordingly, we investigated the expression levels of STRN1, STRN3, and striatin-4 (STRN4) during key steps of peripheral nervous system development. Interestingly, the expression of all three striatin proteins followed a very similar pattern in wild-type (WT) peripheral nerves, with high levels of expression early in development, when both radial sorting and myelination are ongoing, and a large, persistent reduction in protein levels after these processes are complete at P20 (Fig. 3A-B). This suggests that all striatin proteins may have a role in peripheral nerve development and SC biology. In addition, we found that at P10, a time point during which WT mice should be completing radial sorting but are still actively myelinating axons, there is a significant upregulation of STRN4 in sciatic nerves of Strn3^SCKO^ mice (Fig. 3C-D), suggesting that other striatin proteins may be upregulated to compensate for the loss of STRN3 in SCs during development.

**Figure 3:**
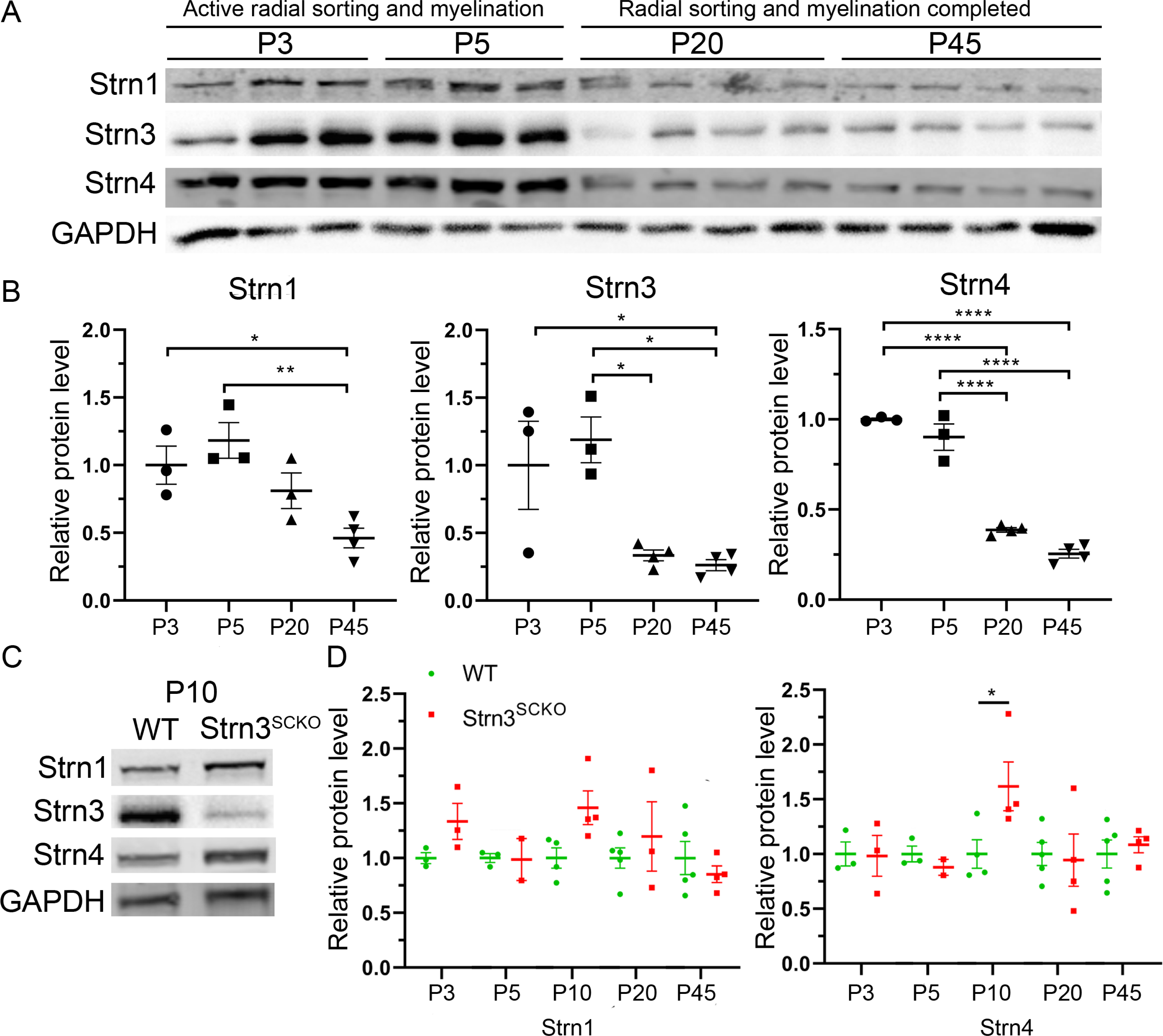
Striatin protein levels are developmentally regulated. (**A**) STRN1, STRN3, and STRN4 protein levels from WT mouse sciatic nerves at P3, P5, P20, and P45. (**B**) Densitometry analysis shows a progressive reduction in STRN1, STRN3, and STRN4 during development in peripheral nerves of WT mice. *N* = 3-4 animals per age group. One-way multiple comparisons ANOVA with Bonferroni post hoc test. STRN1 (*F* (3,9) = 7.66, *p* = 0.0072, *p*_P3-P5_ > 0.9999, *p*_P3-P20_ > 0.9999, *p*_P3-P45_ = 0.0474, *p*_P5-P20_ = 0.3379, *p*_P5-P45_ = 0.0083, *p*_P20-P45_ = 0.3277). STRN3 (*F* (3,10) = 8.772, *p* = 0.0038, *p*_P3-P5_ > 0.9999, *p*_P3-P20_ = 0.0813, *p*_P3-P45_ = 0.0471, *p*_P5-P20_ = 0.0196, *p*_P5-P45_ = 0.0117, *p*_P20-P45_ > 0.9999). STRN4 (*F* (3,10) = 113.4, *p* < 0.0001, *p*_P3-P5_ = 0.5436, *p*_P3-P20_ < 0.0001, *p*_P3-P45_ < 0.0001, *p*_P5-P20_ < 0.0001, *p*_P5-P45_ < 0.0001, *p*_P20-P45_ = 0.0939). (**C**) Representative western blots of STRN1, STRN3, and STRN4 from sciatic nerves of WT and Strn3^SCKO^ mice at P10. (**D**) Densitometry analysis shows an increase in Strn4 at P10 in peripheral nerves of Strn3^SCKO^ mice. *N* = 3-5 animals per genotype per age group. Two-way multiple comparisons ANOVA with Bonferroni post hoc test. STRN1 (*F* (4, 15) age = 1.335; *p* = 0.3026; *F* (1, 11) genotype = 3.755, *p* = 0.0787; *p*_P3_ = 0.5333, *p*_P5_ > 0.9999, *p*_P10_ = 0.0896, *p*_P20_ > 0.9999, *p*_P45_ > 0.9999). STRN4 (*F* age (4, 15) = 1.475, *p* = 0.2591; *F* (1, 12) genotype = 1.305, *p* = 0.2755; *p*_P3_ > 0.9999, *p*_P5_ > 0.9999, *p*_P10_ = 0.0190, *p*_P20_ > 0.9999, *p*_P45_ > 0.9999). Error bars indicate S.E.M. * *p* < 0.05, ** *p* < 0.01, **** *p* < 0.0001.

### Striatins are essential for SC radial sorting

To determine if ablation of *Strn3* in SCs causes radial sorting defects, we analyzed electron micrographs and semithin sections of animals lacking one or two members of the striatin family. We found that loss of any one striatin protein (Strn1^SCKO^, Strn3^SCKO^, or Strn4^SCKO^) resulted in no major defects in the number or density of myelinated fibers or in sciatic nerve thickness at P20 (Fig. 4C). We investigated Strn3^SCKO^ mice more extensively, and while we identified some very mild radial sorting and myelination defects, myelin thickness was not affected (Supplementary Fig. 6A-B). However, ablation of either *Strn1* or *Strn4* in combination with *Strn3* to produce double SCKO mice (Strn1/3^dSCKO^ and Strn3/4^dSCKO^) resulted in a dramatic reduction in the total number and density of myelinated fibers, the presence of immature bundles of unsorted axons, aberrantly myelinated axons inside unsorted bundles, sorted but amyelinated axons, and thinner, hypoplastic sciatic nerves at P20 (Fig. 4). Strn1/3^dSCKO^ mice also retained thinner myelin as late as P60, the latest time point analyzed, suggesting a persistent myelination defect even in SCs that do establish a 1:1 relationship with axons (Supplementary Fig. 7). Strn1/3^dSCKO^ mice were not maintained beyond P60 due to severe, progressive hindlimb muscle wasting and paralysis that begins to manifest shortly after P20. While both Strn1/3^dSCKO^ and Strn3/4^dSCKO^ mice had many massive bundles of unsorted, mixed-caliber axons throughout the sciatic nerve (Fig. 4), it was Strn1/3^dSCKO^ mice that had the most severe axonal sorting phenotype, which persisted as late as P60. An apparent, though much less severe, axonal sorting phenotype was also observed in sciatic nerves of Strn1/4^dSCKO^ mice (Fig. 4). Together, these data suggest that the striatins are essential for SC radial sorting and subsequent myelination during peripheral nerve development, and that the individual striatin protein family members can partially compensate for each other’s function.

**Figure 4:**
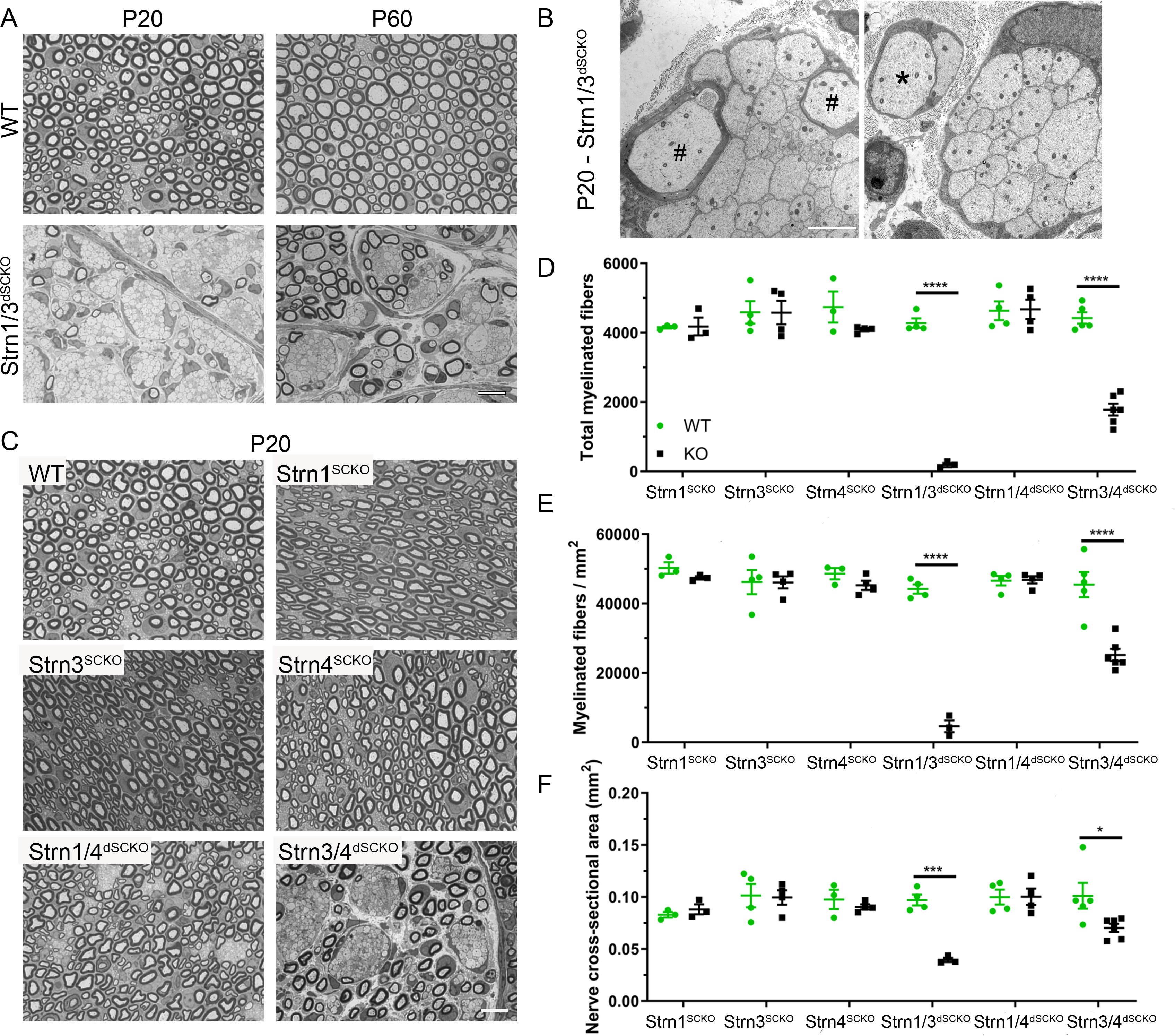
Loss of striatin proteins in SCs results in mild to severe developmental defects. (**A**-**C**) Semithin sections and (**B**) electron micrographs of WT and striatin mutant sciatic nerves at P20 (**A**, **B, C**) and P60 (**A**). At P20, Strn1/3^dSCKO^ present with immature bundles of unsorted axons, myelinated axons inside unsorted bundles (#), and amyelinated axons (*). Scale bar = 10 μm (**A, C**), 2 μm (**B**). (**D-F**) At P20, sciatic nerves of Strn1/3^dSCKO^ and Strn3/4^dSCKO^ mice have significantly fewer total myelinated fibers, a lower density of myelinated fibers, and thinner, hypoplastic sciatic nerves. *N* = 3-6 animals per genotype. Two-way multiple comparisons ANOVA with Bonferroni post hoc test. Total fibers (*F* (5, 35) model = 32.90; *p* < 0.0001; *F* (1, 35) WT vs KO = 76.14, *p* < 0.0001; *p*_Strn1-SCKO_ >0.9999, *p*_Strn3-SCKO_ >0.9999, *p*_Strn4-SCKO_ = 0.4678, *p*_Strn1/3-dSCKO_ < 0.0001, *p*_Strn1/4-dSCKO_ >0.9999, *p*_Strn3/4-dSCKO_ < 0.0001). Fiber density (*F* (5, 35) model = 34.94; *p* < 0.0001; *F* (1, 35) WT vs KO = 71.50, *p* < 0.0001; *p*_Strn1-SCKO_ >0.9999, *p*_Strn3-SCKO_ >0.9999, *p*_Strn4-SCKO_ >0.9999, *p*_Strn1/3-dSCKO_ < 0.0001, *p*_Strn1/4-dSCKO_ >0.9999, *p*_Strn3/4-dSCKO_ < 0.0001). Nerve area (*F* (5, 35) model = 4.602; *p* = 0.0025; *F* (1, 35) WT vs KO = 11.41, *p* = 0.0018; *p*_Strn1-SCKO_ >0.9999, *p*_Strn3-SCKO_ >0.9999, *p*_Strn4-SCKO_ >0.9999, *p*_Strn1/3-dSCKO_ = 0.0001, *p*_Strn1/4-dSCKO_ >0.9999, *p*_Strn3/4-dSCKO_ = 0.0117). Error bars indicate S.E.M. * *p* < 0.05, *** *p* < 0.001, **** *p* < 0.0001.

### STRN1 and STRN3 are required for Hippo pathway regulation in SCs

The striatin proteins function as the center scaffolds of STRIPAK complexes and are necessary for linking together many other members of the complex (*Goudreault, et al., 2009*). Consistent with this, we found that deletion of *Strn1* and *Strn3* in SCs alters the protein levels of other STRIPAK complex members, namely MOB4, STRIP1, and CCM3, in both P5 and P20 peripheral nerves (Supplementary Fig. 6). In addition, while we were expecting compensatory upregulation of STRN4 following loss of STRN1 and STRN3, we found that *Strn4* mRNA and protein levels were also reduced in peripheral nerves of Strn1/3^dSCKO^ mice, suggesting a severe impairment of STRIPAK complexes (Supplementary Fig. 8).

One of the major roles for STRIPAK complexes is to negatively regulate the Hippo pathway, which occurs by dephosphorylation and inactivation of MST1 and MST2 by the STRIPAK-associated PP2A phosphatase (*Bae, et al., 2017; Chen, et al., 2019; Gordon, et al., 2011; Jeong, et al., 2021; Seo, et al., 2020; Tang, et al., 2019; Tang, et al., 2020; Zheng, et al., 2017*). Thus, to further explore the mechanisms underlying the developmental defects in peripheral nerves of Strn1/3^dSCKO^ mice, we next asked if the loss of STRN1 and STRN3 and the subsequent reduction in other STRIPAK proteins causes an increase in the phosphorylation of MST1, MST2, YAP, and TAZ. We found that MST1/2 phosphorylation is significantly increased at both P5 and P20 in Strn1/3^dSCKO^ peripheral nerves (Fig. 5C-E). We also observed increased phosphorylation of YAP and TAZ at P20 in Strn1/3^dSCKO^ peripheral nerves (Fig. 5C-E). In addition to these expected results on the phosphorylation of the Hippo pathway, we also found that the absence of STRN1 and STRN3 in SCs led to a decrease in total MST1/2, YAP, and TAZ protein levels at P5 and P20 (Fig. 5C-E). These results were partially correlated to a reduction of *Mst1*, *Mst2*, and *Yap* mRNA levels (Fig. 5A-B). While it is known that phosphorylation of YAP and TAZ leads to their cytoplasmic retention and subsequent degradation, the global reduction of mRNA and protein levels of various members of the Hippo pathway was unexpected. Together, these data suggest that the loss of STRN1 and STRN3 in SCs causes inhibition of the Hippo pathway via the reduction of gene expression of several Hippo pathway members, and through increased phosphorylation of YAP and TAZ, which leads to their cytosolic sequestration and / or degradation.

**Figure 5:**
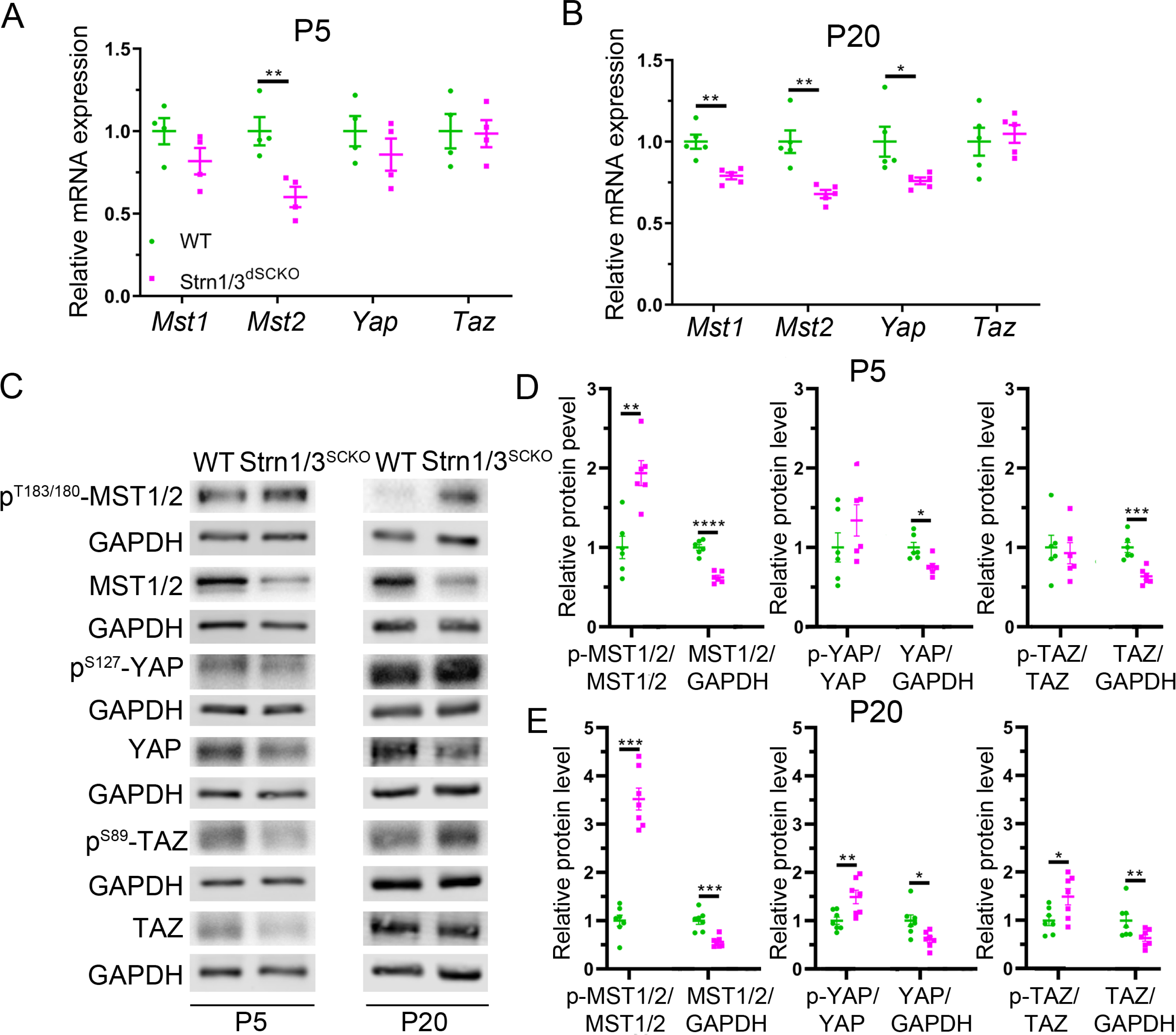
Strn1/3 ablation in SCs results in Hippo pathway dysregulation. (**A-B**) mRNA expression of *Mst1*, *Mst2, Yap, and Taz* from pooled sciatic nerves and brachial plexuses of WT and Strn1/3^dSCKO^ mice at P5 and P20. *Mst2* mRNA level is decreased at P5 and P20, while *Mst1* and *Yap* mRNA levels are decreased at P20 in peripheral nerves of Strn1/3^SCKO^ mice. *N* = 4-5 samples per genotype. P5 samples were pooled from 3-6 animals each. Unpaired two-tailed *t*-test. P5 [*Mst1* (*t* = 1.625, df = 6, *p* = 0.1553), *Mst2* (*t* = 3.789, df = 6, *p* = 0.0091), *Yap* (*t* = 1.061, df = 6, *p* = 0.3296), *Taz* (*t* = 0.1119, df = 6, *p* = 0.9145)]. P20 [*Mst1* (*t* = 4.363, df = 8, *p* = 0.0024), *Mst2* (*t* = 4.368, df = 8, *p* = 0.0024), *Yap* (*t* = 2.562, df = 8, *p* = 0.0335), *Taz* (*t* = 0.4670, df = 8, *p* = 0.6529)]. (**C**) Representative western blots of several Hippo pathways effectors from pooled sciatic nerves and brachial plexuses of WT and Strn1/3^dSCKO^ mice at P5 and P20. (**D-E**) Densitometry analysis shows a reduction in total MST1/2, YAP, and TAZ at P5 and P20, an increase in p-MST1/2 at P5, and an increase in p-MST1/2, p-YAP, and p-TAZ at P20 in peripheral nerves of Strn1/3^dSCKO^ mice. *N* = 6-7 samples per genotype. P5 samples were pooled from 3-6 animals each. Unpaired two-tailed *t*-test. P5 [p-MST1/2 (*t* = 4.431, df = 10, *p* = 0.0013), MST1/2 (*t* = 8.115, df = 10, *p* < 0.0001), p-YAP (*t* = 1.267, df = 10, *p* = 0.2340), YAP (*t* = 3.119, df = 10, *p* = 0.0109), p-TAZ (*t* = 0.3535, df = 10, *p* = 0.7311), TAZ (*t* = 4.803, df = 10, *p* = 0.0007)]. P20 [p-MST1/2 (*t* = 6.906, df = 12, *p* < 0.0001), MST1/2 (*t* = 5.089, df = 12, *p* = 0.0003), p-YAP (*t* = 3.089, df = 12, *p* = 0.0094), YAP (*t* = 2.890, df = 12, *p* = 0.0136), p-TAZ (*t* = 2.534, df = 12, *p* = 0.0267), TAZ (*t* = 2.378, df = 12, *p* = 0.0349)]. Error bars indicate S.E.M. * *p* < 0.05, ** *p* < 0.01, *** *p* < 0.001, **** *p* < 0.0001.

### Rac1 is required for Hippo pathway regulation in SCs

We next sought to clarify if, and how, Hippo pathway dysfunction in Strn1/3^dSCKO^ mice is related to its interaction with Rac1. First, we found that there were no differences in the activity of Rac1 or CDC42, a related Rho GTPase, or in the phosphorylation of the Rac1 effectors PAK1 or NF2 at P20 in peripheral nerves of Strn1/3^dSCKO^ mice (Fig. 6). Thus, our data suggest that STRN1 and STRN3 are downstream of Rac1 activity. Second, we looked at the mRNA, total protein levels, and phosphorylation of MST1, MST2, YAP, and TAZ in mice with SC-specific deletion of *Rac1* (Rac1^SCKO^) mice. Overall, our data show that the absence of Rac1 in SCs also leads to a global increase in phosphorylation of Hippo pathway members (Fig. 7C-E). We also found that MST1/2 total protein levels were reduced in Rac1^SCKO^ peripheral nerves, which correlated with reduced *Mst2* mRNA at P5 (Fig. 7A-B). However, unlike Strn1/3^dSCKO^ mice, we did not find altered levels of total YAP or TAZ (Fig. 7C-E). Thus, these data suggest that the loss of Rac1 in SCs causes Hippo pathway overactivation at the level of MST1 and MST2, with subsequent phosphorylation and inhibition of YAP and TAZ.

**Figure 6:**
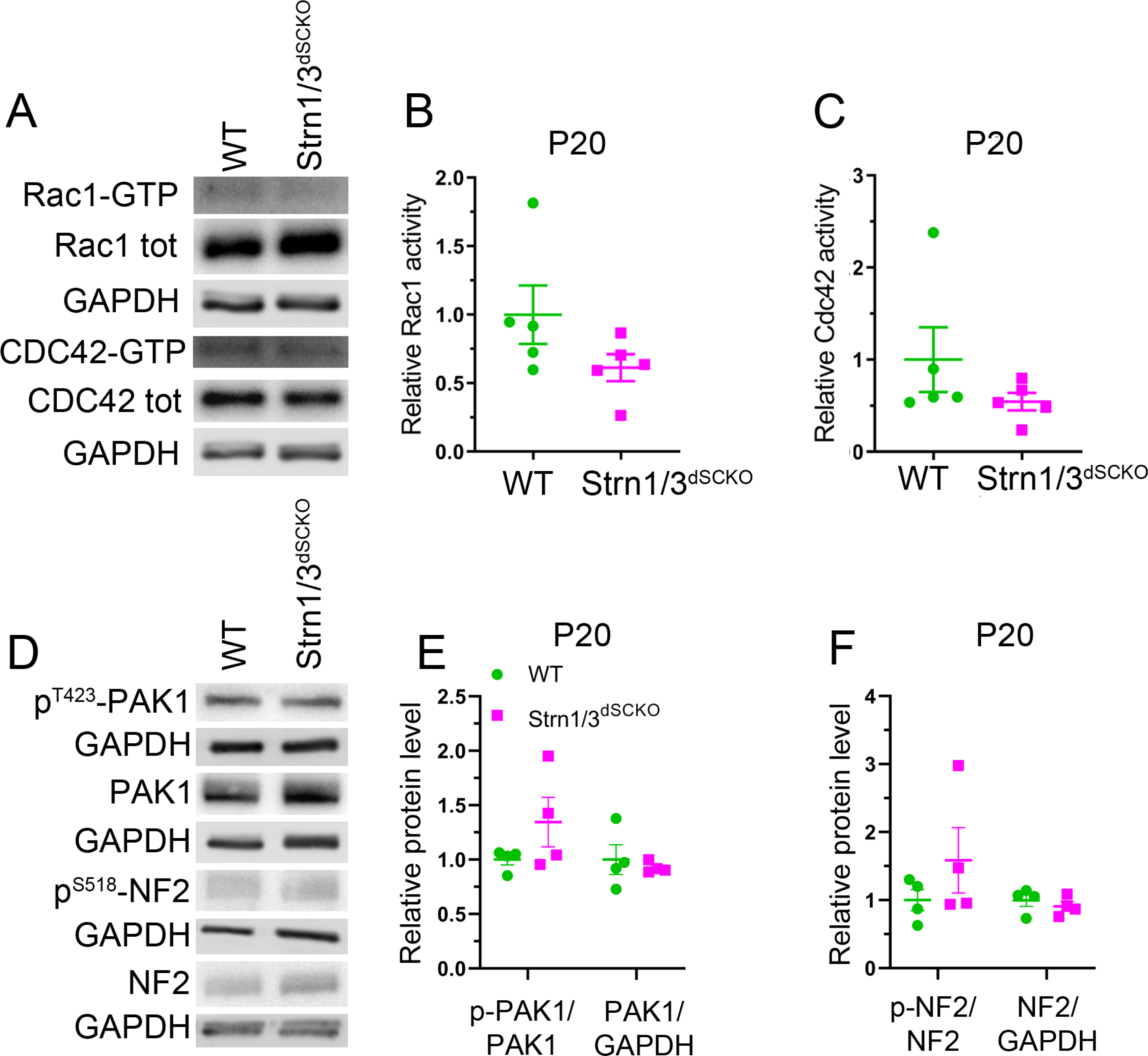
Ablation of *Strn1* and *Strn3* in SCs does not alter Rac1 activity. (**A**) Representative western blots of Rac1 and CDC42 active (GTP) and total (tot) forms from pooled P20 sciatic nerves and brachial plexuses of WT and Strn1/3^dSCKO^ mice. (**B**-**C**) Densitometry analysis shows no significant change in Rac1 or CDC42 activity in peripheral nerves of Strn1/3^dSCKO^ mice at P20. *N* = 5 samples per genotype. Samples were pooled from 3-5 animals each. Unpaired two-tailed *t*-test. Rac1 activity (*t* = 1.649, df = 8, *p* = 0.1379), CDC42 activity (*t* = 1.257, df = 8, *p* = 0.2443). (**D**) Representative western blots of PAK1 and NF2 phosphorylated and total forms from P20 sciatic nerves and brachial plexuses of WT and Strn1/3^dSCKO^ mice. (**E**-**F**) Densitometry analysis shows no significant change in phosphorylation or total levels of the Rac1/CDC42 effectors PAK1 or NF2 in peripheral nerves of Strn1/3^dSCKO^ mice at P20. *N* = 4 animals per genotype. Unpaired two-tailed *t*-test. p-PAK1 (*t* = 1.484, df = 6, *p* = 0.1884), PAK1 (*t* = 0.5307, df = 6, *p* = 0.6147), p-NF2 (*t* = 1.160, df = 6, *p* = 0.2900), NF2 (*t* = 0.7988, df = 6, *p* = 0.4548). Error bars indicate S.E.M.

**Figure 7:**
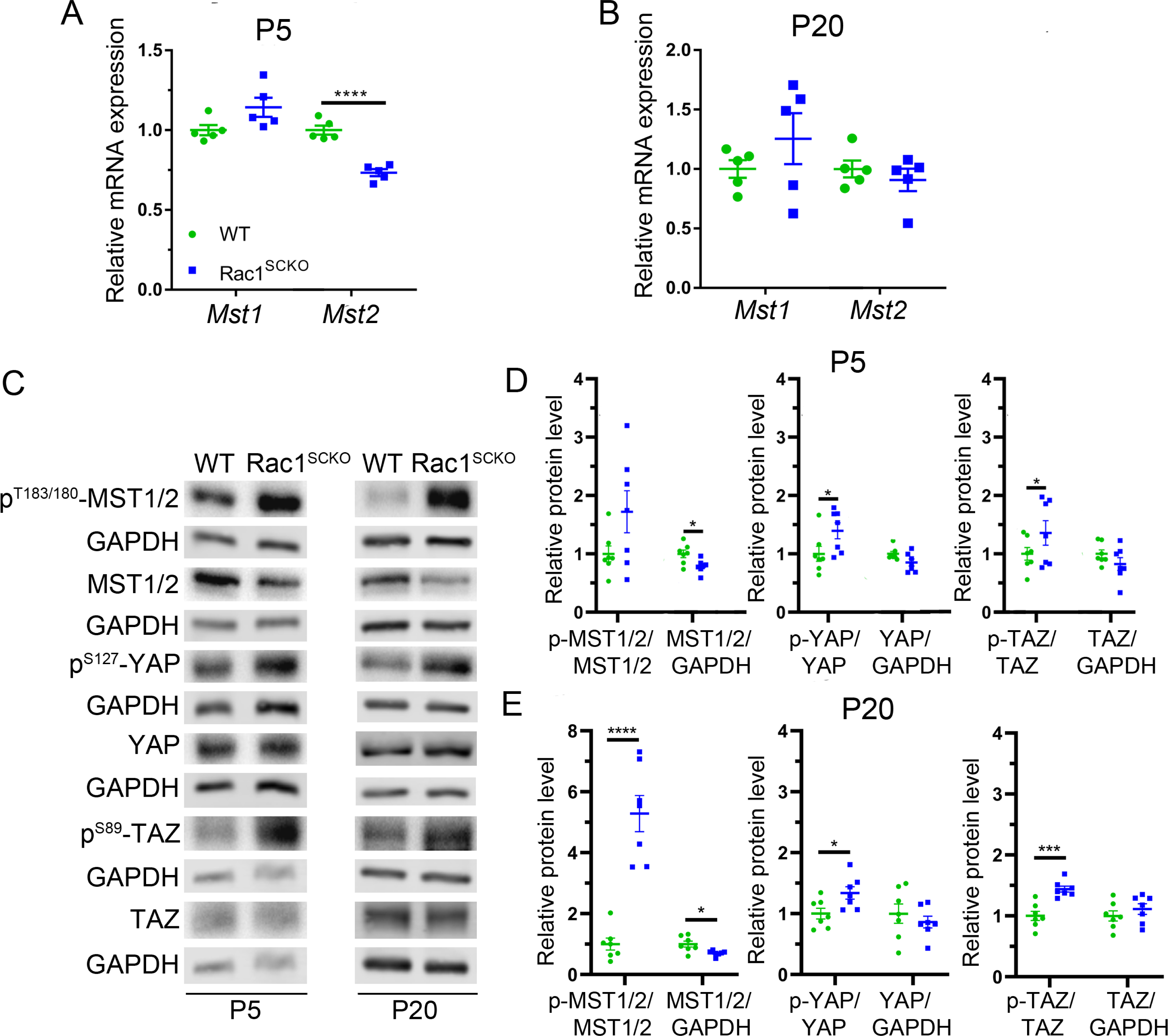
*Rac1* ablation in SCs results in Hippo pathway dysregulation. (**A-B**) mRNA expression of *Mst1* and *Mst2* from pooled sciatic nerves and brachial plexuses of WT and Rac1^SCKO^ mice at P5 and P20. *Mst2* mRNA level is decreased at P5 in peripheral nerves of Rac1^SCKO^ mice. *N* = 5 samples per genotype. P5 samples were pooled from 3-6 animals each. Unpaired two-tailed *t*-test. P5 [*Mst1* (*t* = 2.105, df = 8, *p* = 0.0684), *Mst2* (*t* = 7.480, df = 8, *p* < 0.0001)]. P20 [*Mst1* (*t* = 1.123, df = 8, *p* = 0.2942), *Mst2* (*t* = 0.7793, df = 8, *p* = 0.4582)]. (**C**) Representative western blots of several Hippo pathways members from pooled sciatic nerves and brachial plexuses of WT and Rac1^SCKO^ mice at P5 and P20. (**D-E**) Densitometry analysis shows a reduction in MST1/2 total protein levels at P5 and P20, an increase in p-MST1/2 at P20, and an increase in p-YAP and p-TAZ at P5 and P20 in peripheral nerves of Rac1^SCKO^ mice. *N* = 6-7 samples per genotype. P5 samples were pooled from 3-6 animals each. Unpaired two-tailed *t*-test. P5 [p-MST1/2 (*t* = 1.965, df = 12, *p* = 0.0730), MST1/2 (*t* = 2.750, df = 12, *p* = 0.0176), p-YAP (*t* = 2.998, df = 12, *p* = 0.0111), YAP (*t* = 2.059, df = 12, *p* = 0.0619), p-TAZ (*t* = 2.794, df = 12, *p* = 0.0162), TAZ (*t* = 1.334, df = 12, *p* = 0.2069)]. P20 [p-MST1/2 (*t* = 6.877, df = 12, *p* < 0.0001), MST1/2 (*t* = 2.908, df = 12, *p* = 0.0131), p-YAP (*t* = 2.512, df = 12, *p* = 0.0273), YAP (*t* = 0.7448, df = 12, *p* = 0.4707), p-TAZ (*t* = 4.867, df = 12, *p* = 0.0004), TAZ (*t* = 0.9376, df = 12, *p* = 0.3670)]. Error bars indicate S.E.M. \**p* < 0.05, ** *p* < 0.01, *** *p* < 0.001, **** *p* < 0.0001.

### STRN1, STRN3, and Rac1 regulate SC laminin receptor expression and differentiation

Multiple studies have found that SC expression of several laminin receptors is dependent on YAP and TAZ-mediated transcription (*Deng, et al., 2017; Poitelon, et al., 2016*). In addition, YAP and TAZ-mediated transcription is necessary for SC expression of the master transcription factor *Egr2* (*Deng, et al., 2017; Grove, et al., 2017; Poitelon, et al., 2016*). We therefore hypothesized that the peripheral nerve developmental defects observed in Strn1/3^dSCKO^ mice may be associated with a reduction in the YAP/TAZ-dependent expression of laminin receptors and *Egr2*, with a subsequent retention of *Oct6* expression and failure of differentiation (*Decker, et al., 2006; Ghislain and Charnay, 2006; Le, et al., 2005a; Le, et al., 2005b; Mager, et al., 2008; Zorick, et al., 1996; Zorick, et al., 1999*).

Gene expression analyses revealed several laminin receptors (*Itga6*, *Itgb1*, and *Itgb4*) that were transcriptionally downregulated in Strn1/3^dSCKO^ peripheral nerves at P5, with downregulation of *Itga6* persisting until P20 (Fig. 8A-B). At the protein level, we found reduced integrin α6, integrin β1, and integrin β4 subunits, as well as β-dystroglycan at both P5 and P20 in peripheral nerves of Strn1/3^dSCKO^ mice, suggesting an early and persistent defect in laminin receptor expression (Fig. 8C-E, Fig. 9). Absence of STRN1 and STRN3 also led to an increase in *Oct6* mRNA and protein levels and a decrease in *Egr2* mRNA and protein levels (Fig. 8). Because we showed in our prior works that defect in radial sorting in YAP and TAZ mutants were associated with defects in Schwann cells proliferation (Poitelon et al 2016), we assessed the number of proliferative Schwann cells. While we did not observe significant reduction in SC number or SC proliferation at P5 (Fig.8F), we found that at P20, the number of proliferative Schwann cells was increased in Strn1/3^dSCKO^, correlating with the increased levels of OCT6, a marker for immature Schwann cells (Fig. 8G). These findings support our observations in Strn1/3^dSCKO^ sciatic nerves (Fig. 4) of a severe radial sorting delay with an associated temporal shift in SC proliferation, excess number of immature SCs (OCT6+) and a reduced number of mature, myelinating SCs (EGR2+).

**Figure 8:**
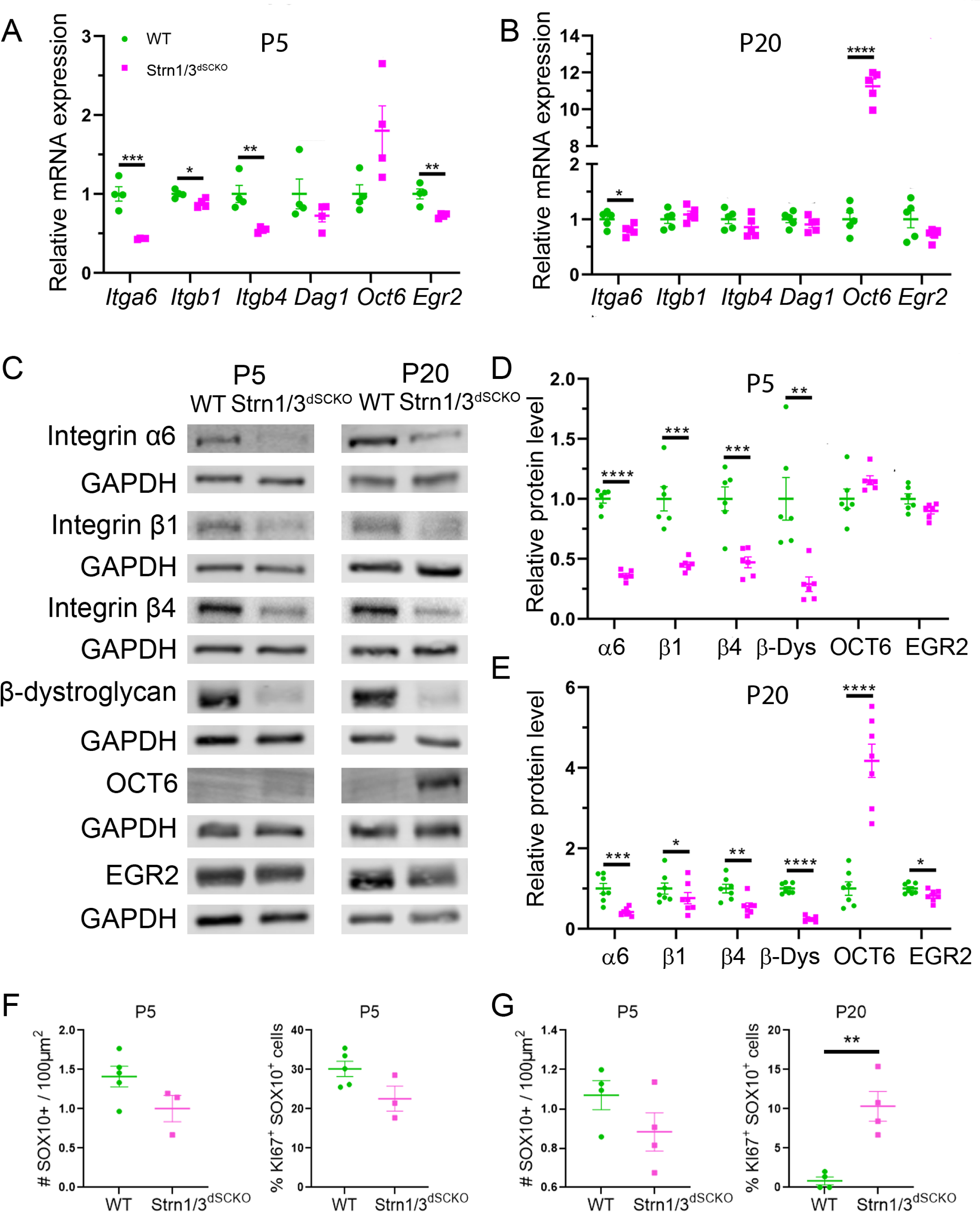
*Strn1* and *Strn3* ablation in SCs results in dysregulated mRNA and protein expression of laminin receptors and SC transcriptional regulators. (**A**-**B**) mRNA expression of several laminin receptors (encoding for integrin subunit α6 (*Itga6*), β1 (*Itgb1*), β4 (*Itgb4*), and β-dystroglycan (*Dag1*)) and SC development transcriptional regulators (*Oct6* and *Egr2*) from pooled sciatic nerves and brachial plexuses of WT and Strn1/3^dSCKO^ mice at P5 and P20. *Itgb1, Itgtb4* and *Egr2* mRNA levels are decreased at P5, *Itga6* mRNA level is decreased at P5 and P20, and *Oct6* mRNA level is increased at P20 in peripheral nerves of Strn1/3^dSCKO^ mice. *N* = 4-5 samples per genotype. P5 samples were pooled from 3-6 animals each. Unpaired two-tailed *t*-test. P5 [*Itga6* (*t* = 6.236, df = 6, *p* = 0.0008), *Itgb1* (*t* = 3.094, df = 6, *p* = 0.0213), *Itgb4* (*t* = 4.102, df = 6, *p* = 0.0063), *Dag1* (*t* = 1.341, df = 6, *p* = 0.2284), *Oct6* (*t* = 2.380, df = 6, *p* = 0.0548), *Egr2* (*t* = 4.288, df = 6, *p* = 0.0052)]. P20 [*Itga6* (*t* = 2.487, df = 8, *p* = 0.0377), *Itgb1* (*t* = 0.9136, df = 8, *p* = 0.3876), *Itgb4* (*t* = 1.254, df = 8, *p* = 0.2451), *Dag1* (*t* = 1.103, df = 8, *p* = 0.3020), *Oct6* (*t* = 25.75, df = 8, *p* < 0.0001), *Egr2* (*t* = 1.722, df = 8, *p* = 0.1233)]. (**C**) Representative western blots of several laminin receptors and SC development transcriptional regulators from pooled sciatic nerves and brachial plexuses of WT and Strn1/3^dSCKO^ mice at P5 and P20. (**D**-**E**) Densitometry analysis shows a reduction in integrin α6, β1, and β4 subunits and β-dystroglycan at P5 and P20, an increase in OCT6 at P20, and a reduction in EGR2 at P20 in peripheral nerves of Strn1/3^dSCKO^ mice. *N* = 6-7 samples per genotype. P5 samples were pooled from 3-6 animals each. Unpaired two-tailed *t*-test. P5 [integrin α6 (*t* = 16.27, df = 10, *p* < 0.0001), integrin β1 (*t* = 5.328, df = 10, *p* = 0.0003), integrin β4 (*t* = 4.826, df = 10, *p* = 0.0007), β-dystroglycan (*t* = 3.777, df = 10, *p* = 0.0036), OCT6 (*t* = 1.717, df = 10, *p* = 0.1168), EGR2 (*t* = 2.053, df = 10, *p* = 0.0671)]. P20 [integrin α6 (*t* = 4.446, df = 12, *p* = 0.0008), integrin β1 (*t* = 2.994, df = 12, *p* = 0.0112), integrin β4 (*t* = 3.277, df = 12, *p* = 0.0066), β-dystroglycan (*t* = 13.65, df = 12, *p* < 0.0001), OCT6 (*t* = 7.137, df = 12, *p* < 0.0001), EGR2 (*t* = 2.443, df = 12, *p* = 0.0310)]. (**F-G**) Quantification of the total number of Schwann cells (SOX10+) and proliferative Schwann cells (SOX10+ KI67+) at P5 and P20 show an increase in Schwann cells proliferation at P20 in peripheral nerves of Strn1/3^dSCKO^ mice. *N* = 5-3 samples per genotype. Unpaired two-tailed *t*-test. P5 [SOX10+ (*t* = 1.908, df = 6, *p* = 0.1051), KI67+SOX10+ (*t* = 2.179, df = 6, *p* = 0.0722). P20 [SOX10+ (*t* = 1.532, df = 6, *p* = 0.1764), KI67+SOX10+ (*t* = 4.847, df = 6, *p* = 0.0029). Error bars indicate S.E.M. \**p* < 0.05, ** *p* < 0.01, *** *p* < 0.001, **** *p* < 0.0001.

**Figure 9:**
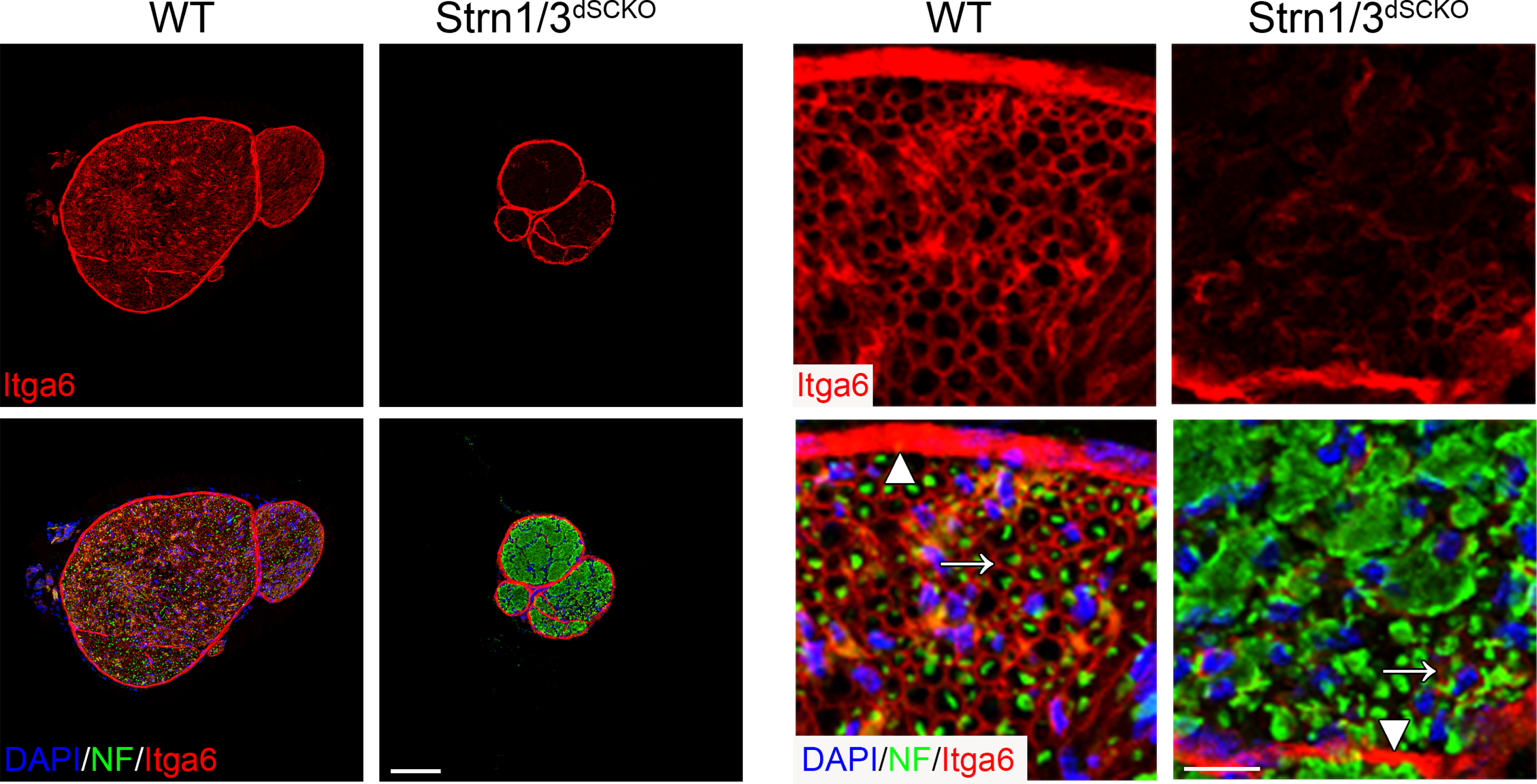
*Strn1* and *Strn3* ablation in SCs results in reduced integrin α6 subunit expression. Sciatic nerve cross sections from WT and Strn1/3^dSCKO^ mice at P20 stained for integrin α6 subunit (Itga6, red). Axons are labeled by neurofascin (NF, green) and nuclei are labeled with DAPI (blue). White arrows indicate integrin α6 expression by SCs surrounding axons and white arrowheads indicate integrin α6 expression by perineurial cells in the perineurium. Integrin α6 appears drastically reduced in SCs, but not in the perineurium, in sciatic nerves of Strn1/3^dSCKO^ mice at P20. Scale bar = 100µm (left), 10µm (right).

Finally, we found that Rac1^SCKO^ peripheral nerves recapitulate many of the phenotypes observed in Strn1/3^dSCKO^ mice on laminin receptor, *Oct6*, and *Egr2* mRNA and protein levels (Fig. 10). These data suggest that the absence of Rac1 in SCs is associated with a downregulation of laminin receptors and EGR2 with a concomitant upregulation of OCT6. Altogether, our results strongly indicate that Rac1, STRN1, and STRN3 are modulating developmentally crucial gene expression in SCs through the Hippo pathway, YAP, and TAZ.

**Figure 10:**
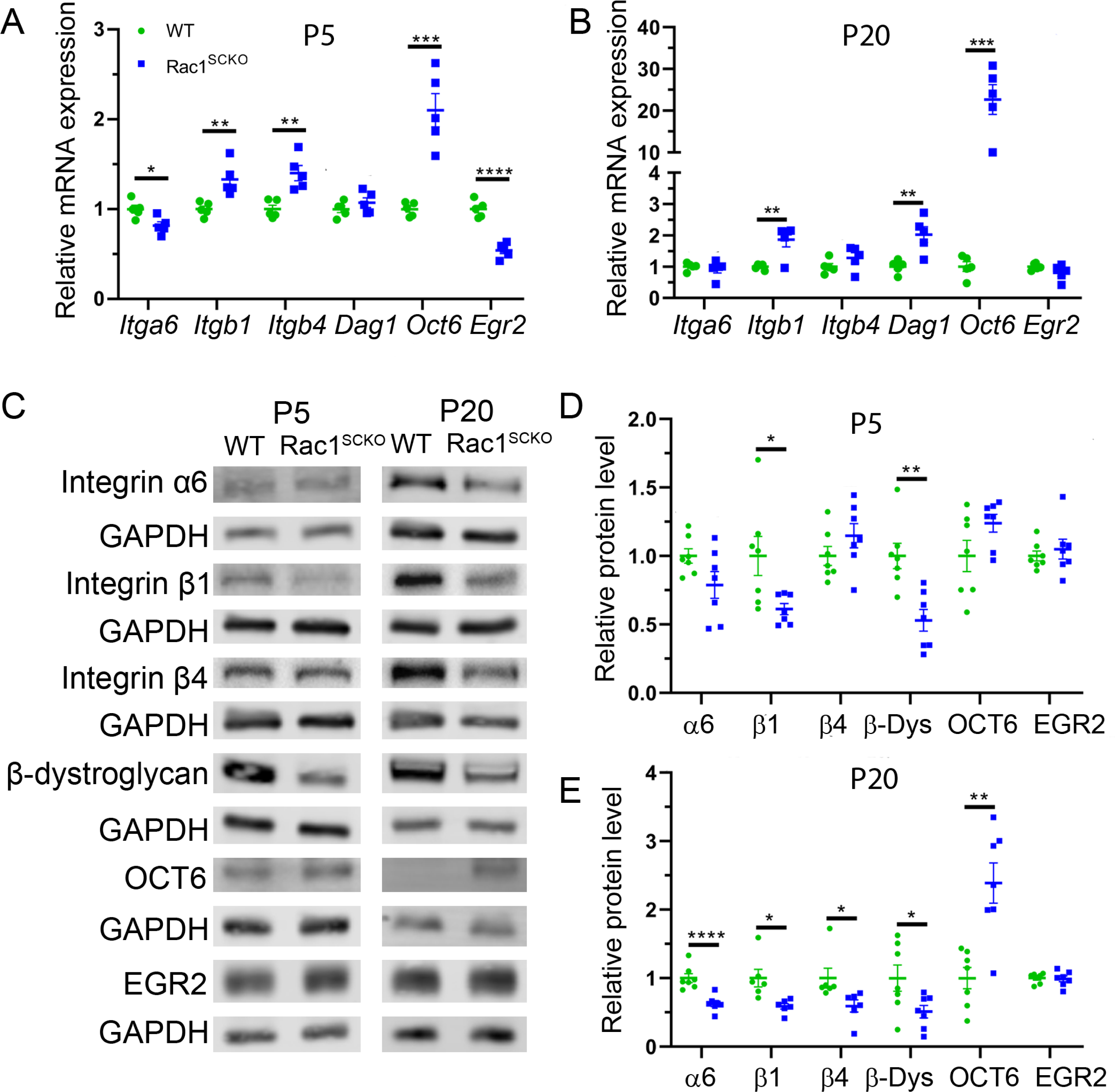
*Rac1* ablation in SCs results in dysregulated mRNA and protein expression of laminin receptors and SC transcriptional regulators. (**A**-**B**) mRNA expression of several laminin receptors (encoding for integrin subunit α6 (*Itga6*), β1 (*Itgb1*), β4 (*Itgb4*), and β-dystroglycan (*Dag1*)) and SC development transcriptional regulators (*Oct6* and *Egr2*) from pooled sciatic nerves and brachial plexuses of WT and Rac1^SCKO^ mice at P5 and P20. *Itga6* and *Egr2* mRNA levels are decreased at P5, *Itgb1* and *Oct6* mRNA levels are increased at P5 and P20, *Itgb4* mRNA level is increased at P5, and *Dag1* mRNA level is increased at P20 in peripheral nerves of Rac1^SCKO^ mice. *N* = 5 samples per genotype. P5 samples were pooled from 3-6 animals each. Unpaired two-tailed *t*-test. P5 [*Itga6* (*t* = 3.042, df = 8, *p* = 0.0160), *Itgb1* (*t* = 3.771, df = 8, *p* = 0.0055), *Itgb4* (*t* = 4.199, df = 8, *p* = 0.0030), *Dag1* (*t* = 1.086, df = 8, *p* = 0.3089), *Oct6* (*t* = 5.829, df = 8, *p* = 0.0004), *Egr2* (*t* = 8.147, df = 8, *p* < 0.0001)]. P20 [*Itga6* (*t* = 0.5845, df = 8, *p* = 0.5750), *Itgb1* (*t* = 3.717, df = 8, *p* = 0.0059), *Itgb4* (*t* = 1.418, df = 8, *p* = 0.1940), *Dag1* (*t* = 3.793, df = 8, *p* = 0.0053), *Oct6* (*t* = 6.063, df = 8, *p* = 0.0003), *Egr2* (*t* = 1.580, df = 8, *p* = 0.1529)]. (**C**) Representative western blots of several laminin receptors and SC development transcriptional regulators from pooled sciatic nerves and brachial plexuses of WT and Rac1^SCKO^ mice at P5 and P20. (**D**-**E**) Densitometry analysis shows a reduction in integrin β1 subunit and β-dystroglycan (β-Dys) at P5 and P20, integrin α6 and β4 subunits at P20, and an increase in OCT6 at P20 in peripheral nerves of Rac1^SCKO^ mice. *N* = 6-7 samples per genotype. P5 samples were pooled from 3-6 animals each. Unpaired two-tailed *t*-test. P5 [integrin α6 (*t* = 0.2671, df = 12, *p* = 0.7939), integrin β1 (*t* = 2.619, df = 12, *p* = 0.0224), integrin β4 (*t* = 1.313, df = 12, *p* = 0.2139), β-dystroglycan (*t* = 4.697, df = 12, *p* = 0.0005), OCT6 (*t* = 4.127, df = 12, *p* = 0.0014), EGR2 (*t* = 0.6095, df = 12, *p* = 0.5535)]. P20 [integrin α6 (*t* = 4.791, df = 12, *p* = 0.0004), integrin β1 (*t* = 3.086, df = 10, *p* = 0.0115), integrin β4 (*t* = 2.389, df = 10, *p* = 0.0380), β-dystroglycan (*t* = 2.304, df = 12, *p* = 0.0399), OCT6 (*t* = 4.166, df = 12, *p* = 0.0013), EGR2 (*t* = 0.2044, df = 12, *p* = 0.8415)]. Error bars indicate S.E.M. \**p* < 0.05, ** *p* < 0.01, *** *p* < 0.001, **** *p* < 0.0001.

## DISCUSSION

In this study, we identified a previously unknown interaction between Rac1 and two STRIPAK complex members, STRN3 and MOB4. We found that members of the striatin family of proteins are essential in SCs for axonal radial sorting and SC differentiation. Additionally, loss of Rac1 or STRN1/3 resulted in dysregulation of Hippo pathway signaling, YAP/TAZ activity, laminin receptor expression, and expression of the master transcription factor EGR2. Furthermore, we found that the loss of STRN1 and STRN3 resulted in a global breakdown in the abundance of other STRIPAK complex members.

Signals from the SC-derived basal lamina, transmitted via laminin receptors, are required for SC proliferation, survival, differentiation, apical-basal polarity, radial sorting, initiation of myelination, radial and longitudinal growth of the myelin sheath, and nodal clustering of voltage-gated sodium channels (*Berti, et al., 2011; Court, et al., 2009; Feltri, et al., 2002; Feltri, et al., 2016; Nodari, et al., 2008; Nodari, et al., 2007; Occhi, et al., 2005; Pellegatta, et al., 2013; Previtali, et al., 2003a; Previtali, et al., 2003b; Saito, et al., 2003; Yu, et al., 2005*). Among the signals initiated upon receptor-ECM ligand engagement is Rho GTPase activity. Rac1 and a closely related Rho GTPase, CDC42, have previously been shown to be required for SCs to radially sort and myelinate axons (*Benninger, et al., 2007; Guo, et al., 2012; Guo, et al., 2013; Nodari, et al., 2007*). Rac1 has also been shown to be required for the early SC response to peripheral nerve injury, an organized process by which SCs initiate myelin fragmentation and digestion (*Jung, et al., 2011*). Expression of dominant negative NF2, a downstream Rac1/CDC42 effector, partially rescues the developmental myelination defect in both Rac1 and CDC42 mutant mice, but only rescues the radial sorting defect in CDC42 mutant mice (*Guo, et al., 2012; Guo, et al., 2013*). Our understanding of how, precisely, NF2 mediates this partial rescue, the difference between Rac1 and CDC42 mutant mice, and the additional mechanisms mediating radial sorting and myelination defects in these models remains limited.

Previous studies have found conflicting reports about the influence of Rac1 on Hippo pathway signaling and YAP/TAZ activity in a variety of cell types in vitro (*Esposito, et al., 2022; Sabra, et al., 2017; Talwar, et al., 2021; Zhao, et al., 2012*). Additionally, multiple groups have reported mechanisms by which NF2, a Rac1 effector, influences STRIPAK-independent and STRIPAK-dependent regulation of the Hippo pathway (*Bae, et al., 2017; Chen, et al., 2019; Li, et al., 2015; Sabra, et al., 2017; Yin, et al., 2013*). One STRIPAK-independent mechanism suggests that NF2 directly interacts with AMOT, a YAP effector and binding partner, thereby acting as a bridge to promote LATS1/2-mediated phosphorylation of YAP (*Sabra, et al., 2017*). A STRIPAK-dependent mechanism suggests that SAV1, an MST1/2 adaptor protein, preserves MST1/2 activity by antagonizing STRIPAK/PP2A-mediated dephosphorylation of p-MST1/2 (Thr183/180) (*Bae, et al., 2017*). Deletion of the N-terminal portion of SAV1, which includes its NF2 binding domain, reduced its ability to preserve MST1/2 activity, suggesting that NF2 may influence SAV1-mediated antagonism of STRIPAK. Another STRIPAK-dependent mechanism suggests that NF2 interacts with STRIPAK via the STRIP1 STRIPAK adaptor protein to prevent association of the PP2A phosphatase with STRIPAK complexes (*Chen, et al., 2019*). Of note, loss of NF2 in SCs has been demonstrated to result in defects in axonal regeneration and remyelination following peripheral nerve injury, which is rescued by co-deletion of *Yap* in SCs (*Mindos, et al., 2017*). However, this result contrasts somewhat with more recent studies that demonstrate a defect in remyelination following peripheral nerve injury in mice lacking both YAP and TAZ in SCs (*Grove, et al., 2020; Jeanette, et al., 2021*). Our results indicate that ablation of *Rac1* in SCs in vivo results in Hippo pathway activation and YAP/TAZ inhibition with a subsequent loss of YAP/TAZ-mediated laminin receptor and transcription factor expression (*Deng, et al., 2017; Grove, et al., 2017; Poitelon, et al., 2016*). We propose that this overactive Hippo signaling, and its downstream effects are responsible for a portion of both the NF2-dependent and NF2-independent peripheral nerve developmental defects in Rac1^SCKO^ and possibly CDC42^SCKO^ mice. We believe that a similar mechanism is responsible for the more severe phenotype observed in Strn1/3^dSCKO^ mice. Further studies could confirm the specific roles of Hippo pathway signaling in these models by targeting the upstream Hippo pathway pharmacologically, such as with the MST1/2 inhibitor XMU-MP-1, or with conditional mutagenesis approaches (*Fan, et al., 2016; Lu, et al., 2010; Wu, et al., 2018*).

Importantly, aside from its function in modulating Hippo pathway signaling, the STRIPAK complex has been implicated in many other cellular functions, as has been extensively reviewed elsewhere (Hwang et al., 2013; Kuck et al., 2019; Kuck et al., 2021). For example, there is evidence for a direct role for STRIPAK in TORC2, JNK, p38 MAPK, and RhoA signaling and in processes such as synaptic terminal formation, mitophagy, and autophagosome fusion (Sakuma et al., 2016; Innokentev et al., 2020; Guo et al., 2023). Thus, it is possible that the phenotype observed in the Strn1/3^dSCKO^ model, is caused by Hippo-independent mechanisms. First, in the Strn1/3^dSCKO^ model, there is a severe disruption in the expression of various laminin receptor subunits, which we believe may alone be sufficient to elicit the extreme morphological phenotype observed (Nodari et al. 2016). Second, it is plausible that there are also direct, STRIPAK-mediated effects on the actin cytoskeleton. It has already been demonstrated that Rac1 is essential for lamellipodia formation, process elongation, and injury-induced actin polymerization (Nodari et al., 2007; Benninger et al., 2007; Jung et al., 2011). In addition, Rac1 activity has previously been demonstrated to have multiple pronounced effects on the actin dynamics of peripheral ensheathing glia of Drosophila melanogaster (Sepp et al., 2003). Yet these mechanisms are not mutually exclusive to the previously published literature modulation of the actin cytoskeleton is also known to regulate YAP and TAZ in Hippo-independent manner (Dupont et al. 2011, Aragona et al. 2013).

Our finding of a direct interaction between Rac1 and STRN3/MOB4 suggests a potentially novel mechanism for linking Rac1 activity and Hippo pathway signaling. This is an intriguing possibility, given the diverse roles of YAP/TAZ in SCs and the druggable target nature of Rac1 and various STRIPAK complex subunits. Loss of either Rac1 or STRN1/3 in SCs results in MST1/2 overactivation and YAP/TAZ inhibition by phosphorylation. However, further work is required to prove a STRIPAK-dependent mechanism of Hippo pathway regulation by Rac1. Future studies could seek to clarify the purpose of these Rac1-STRIPAK interactions, perhaps by transgenic depletion of STRN3, MOB4, or PP2A, pharmacologic modulation of PP2A activity with drugs like okadaic acid, disruption of the Rac1-STRN3 or Rac1-MOB4 interactions, or disruption of the STRN3-PP2A interaction necessary for STRIPAK-mediated inhibition of the Hippo pathway. Exciting new technologies for the latter, such as a STRN3-derived Hippo-activating peptide (SHAP) and improved heterocyclic PP2A activators (iHAPs), have recently been described (*Kurppa and Westermarck, 2020; Tang, et al., 2020*).

Interestingly, we found reduced total levels of the MST1/2 kinases following ablation of Rac1 or STRN1/3, and this appeared to be partially mediated by transcriptional repression. To the best of our knowledge, downregulation of total MST1/2 as a regulatory mechanism has not been previously described in any cell type. We propose that this reduction in total MST1/2 could represent a previously undescribed Hippo pathway regulatory feedback loop. In the context of overactive MST1/2 kinase activity, partial suppression of this aberrant signaling may be achieved via a reduction in total protein levels. While our results indicate that this effect is partly transcriptionally mediated, additional investigations will be required to determine how this transcriptional response is initiated and what other mechanisms may be involved. New developments in targeting MST1/2 are an exciting prospect, given the broad interest in modifying Hippo pathway signaling in fields as diverse as developmental disorders, regenerative medicine, and oncology (*Johnson and Halder, 2014*).

To the best of our knowledge, the reduction in protein levels of various STRIPAK complex subunits (STRN4, MOB4, STRIP1, and CCM3) following loss of the central striatin scaffolding proteins has not been previously described. We had anticipated potential protein instability of STRIPAK subunits following *Strn1/3* deletion; however, the additional reduction in STRN4 levels was surprising due to its upregulation following single *Strn3* ablation in SCs. Given the markedly reduced phenotypic severity of STRN1/4^dSCKO^ mice compared to STRN1/3^dSCKO^ or STRN3/4^dSCKO^ mice, our results indicate that STRN3 is the only striatin protein sufficient to perform the majority of striatin family protein functions during SC development alone. Altogether, these results suggest that STRN3 is particularly important for the stability of STRIPAK complexes and for STRIPAK functions. The role of STRN3 in STRIPAK complex integrity may be of particular interest to groups studying the Hippo pathway in various disease contexts, given the involvement of STRIPAK adaptor proteins such as MOB4, STRIP1, and CCM3 in regulating the Hippo pathway in diseases such as pancreatic cancer, gastric cancer, hepatocellular carcinoma, and breast cancer (*Chen, et al., 2018; Chen, et al., 2019; Sun, et al., 2021; Wang, et al., 2021*). Recent studies have also found STRN3 to be unique among the striatins, such as in its ability to form asymmetric homotetramers with a single catalytic STRIPAK core (*Jeong, et al., 2021*). Additionally, while all three striatin proteins and their inhibitory effect on the Hippo pathway have been correlated with human cancers, STRN3 is cited in a particularly wide array of human cancer subtypes, to include various immortalized cancer cell lines (hepatocellular carcinoma, prostatic adenocarcinoma, medulloblastoma), patient-derived primary cultures and biopsy specimens (gastric cancer, hepatocellular carcinoma, cribriform adenocarcinoma of the salivary gland, breast cancer), and estrogen receptor (ER)-dependent clinical outcomes of breast cancer patients (possibly due to inhibition of ERα by STRN3) (*Li, et al., 2023; Migliavacca, et al., 2022; Owosho, et al., 2023; Tan, et al., 2008; Tang, et al., 2020; Tanti, et al., 2015; Zhu, et al., 2024*). We believe that our findings and others make STRN3 a useful starting candidate in future investigations of STRIPAK biology.

It is inherently difficult to parse out radial sorting and myelination defects, as both developmental processes occur in a similar time window. Animals with radial sorting defects can have concomitant myelination defects, such as excess numbers of promyelinating SCs, abnormal myelin features, and / or hypomyelination (*Nodari, et al., 2007; Poitelon, et al., 2016*). However, mutants with radial sorting defects can also present without myelination defects, such as in *Lama2^−/−^* mice (*Ghidinelli, et al., 2017*). While STRN3^SCKO^ and STRN1/3^dSCKO^ mice present with defects in myelin formation, supporting that striatins are required for the myelination step of SC development, the principal role of striatins appears to be in facilitating radial sorting.

In conclusion, our study reveals that striatin proteins are critical for SC radial sorting and subsequent myelination and that loss of STRN1/3 or Rac1 results in a profound deficit in laminin receptor expression and SC differentiation, possibly via dysregulation of Hippo pathway signaling. The influence of the Hippo pathway and YAP/TAZ in regulating peripheral neuropathy disease-associated gene expression, tumor progression in cancers such as MPNSTs, and remyelination with functional recovery following peripheral nerve injury is gaining increased attention (*Deng, et al., 2017; Feltri, et al., 2021; Grove, et al., 2024; Grove, et al., 2017; Grove, et al., 2020; Jeanette, et al., 2021; Laraba, et al., 2023; Lopez-Anido, et al., 2016; Mindos, et al., 2017; Poitelon, et al., 2016; Wang, et al., 2023; Wu, et al., 2018*). Our study adds to the growing body of knowledge about modulation of the Hippo pathway and YAP/TAZ by identifying potentially novel Hippo/YAP/TAZ regulatory machinery in SCs (*Moore, et al., 2024*). Namely, we have demonstrated marked dysregulation of Hippo signaling and YAP/TAZ activity following *Rac1* or *Strn1/3* ablation in SCs. Additionally, our work proposes multiple potentially novel paradigms for modifying Hippo pathway signaling in SCs, such as by targeting the Rac1-STRN3/MOB4 interactions, MST1/2 protein abundance, or STRIPAK complex integrity. The Rac1-STRN3/MOB4 interactions and loss of STRIPAK complex integrity following deletion of *Strn1/3* also uncover intriguing new aspects of basic STRIPAK biology. We posit that further investigation of STRIPAK complexes will establish if they regulate ECM receptor expression or cellular differentiation in other cell types and lead to a better understanding of their influence in modulating a variety of additional cellular functions in SC. This research could spur the development of novel, STRIPAK-targeting therapeutic approaches with applications in diverse fields such as peripheral nerve development, peripheral neuropathies, peripheral nerve tumors, and peripheral nerve regeneration.

## METHODS

### Animal models

All animal experiments were approved by the Institutional Animal Care and Use Committee (IACUC) of Roswell Park Comprehensive Cancer Center (RPCCC) and the State University of New York (SUNY) University at Buffalo protocols UB1188M and UB1194M. Animals were housed in sex-segregated, individually ventilated cages with a maximum of five adult mice per cage. Food and water were provided ad libitum and rooms were maintained at a 12-hour light / dark cycle. MPZ-Cre, Strn1-floxed, Strn3-floxed, Strn4-floxed, and Rac1-floxed mice kept in a C57BL/6 background strain were used in this study (*Feltri, et al., 1999; Nodari, et al., 2007*). All three striatin lines were generated as part of the UC Davis Knockout Mouse Project (KOMP) (*Skarnes, et al., 2011*). Striatin-1 floxed and striatin-3 floxed mice were obtained from the Mutant Mouse Resources & Research Centers (MMRRC) while striatin-4 floxed mice were obtained from the European Mutant Mouse Archive (EMMA) node at the Institute of Molecular Genetics ASCR. For single Schwann cell knockout (SCKO) animals, mice carrying one or two floxed alleles but no MPZ-Cre were used as controls. For double SCKO (dSCKO) animals, mice carrying three or four floxed alleles but no MPZ-Cre were used as controls. No animals were excluded from this study.

### Rac1 activity assay

For each 400 μg tissue sample, 5 μl of settled (20 μl of slurry) glutathione magnetic agarose beads (Thermo Fisher Scientific 78601) were first washed 3x with 800 μl Rac1 lysis buffer (20 mM Tris-HCl pH 7.5, 150 mM NaCl, 1% Triton X-100, 10 mM MgCl_2_, 10 mM NaF, and 1:100 protease inhibitor cocktail) and then resuspended in 800 μl Rac1 lysis buffer with 10 μg GST-PAK-PBD (Cytoskeleton, Inc. PAK01-A). GST-PAK-PBD was conjugated to beads by rotating for 1 hour at room temperature, followed by washing 3x in 800 μl Rac1 wash buffer and then resuspending in 300 μl cold Rac1 wash buffer on ice. Pulverized sciatic nerves and brachial plexuses were prepared as described in supplemental methods. Pulverized tissue was lysed with Rac1 lysis buffer and centrifuged at 13,200 rcf for 1 minute at 4°C. Protein concentration of the supernatant was quantified by Precision Red assay (Cytoskeleton, Inc. ADV02). For GDP and GTP-charged controls, 100 μg of tissue lysate was incubated with 2 μl 0.5 M EDTA and either 5 μl 20 mM GDP (Sigma-Aldrich G7637) or 5 μl 20 mM GTPγS (Sigma-Aldrich G8634) for 15 minutes at 37°C. GDP/GTP control charge reactions were stopped by adding 6 μl 1M MgCl_2_. To pull down active Rac1, 400 μg of tissue lysate (or 100 μg for GDP/GTP-charged controls) was added to GST-PAK-PBD-conjugated beads. Total volume per sample was brought to 800 μl with cold Rac1 lysis buffer and samples were rotated for 1 hour at 4°C. Lysate was removed and beads were washed with 800 μl with cold Rac1 lysis buffer. Rac1 lysis buffer was removed, samples were centrifuged at 1,000 rcf for 5 seconds at 4°C, and then residual buffer was removed. Samples were eluted by boiling in 40 μl 2x Laemmli buffer for 10 minutes at 95°C. Samples were then analyzed by western blot as described in supplemental methods.

### Rac1 pulldown assay

For each 500 μg tissue sample, 5 μg of GST-Rac1 (Cytoskeleton, Inc. RCG01-C) was charged with GDP or GTPγS by incubating for 20 minutes at 37°C in 100 μl GST pulldown buffer (20 mM Tris-HCl pH 7.4, 150 mM NaCl, 0.1% Triton X-100, 10 mM MgCl_2_, and 1:100 protease inhibitor cocktail) containing 20 mM EDTA and1 mM of either GDP or GTPγS. The charge reaction was stopped by adding an additional 2 μl 1M MgCl_2_. For each sample, 5 ul settled glutathione beads (20 μl bead slurry) were rinsed 3x in 800 μl GST pulldown buffer and then resuspended in 400 μl GST pulldown buffer. To conjugate GST-Rac1-GDP/GTPγS to the beads, each ∼100 ul charge reaction was combined with a 400 μl washed bead suspension and rotated for 1 hour at 4°C, washed 3x with 800 μl cold GST pulldown buffer, and then resuspended in 300 μl GST pulldown buffer. GST-Rac1-GDP/GTPγS-conjugated beads were then combined with 500 μg tissue lysed in Rac1 lysis buffer as described above and rotated for 1 hour at 4°C. Samples were rinsed 3x with 800 μl cold GST pulldown buffer, centrifuged at 1,000 rcf for 5 seconds at 4°C, and then residual buffer was removed. Samples were either eluted by boiling in 40 μl 2x Laemmli buffer for 10 minutes at 95°C for analysis by western blot as described above or frozen and shipped to colleagues for analysis by tandem mass spectroscopy (MS/MS). For MS/MS analysis, samples were digested with trypsin to generate tryptic peptides. The Welch difference was calculated as the log2-fold change in protein enrichment between Rac1-GDP and Rac1-GTP fractions. A false discovery rate (FDR) = 0.5 was used as the cutoff value to declare a protein specifically enriched in either fraction.

### Co-immunoprecipitation (co-IP)

Rat SCs (rSC) were cultured in rSC media (described in supplemental methods) on uncoated tissue culture dishes until approximately 80% confluent (*Catignas, et al., 2021*). Cells were then washed with cold PBS and lysed in ice cold IP lysis buffer (PBS + 1% NP-40 + 1 mM EDTA + 1:100 protease inhibitor cocktail). Lysate protein concentration was quantified by BCA assay and 1 mg of protein was combined with 0.25 μg of either rabbit anti-Rac1 experimental antibody (Thermo Fisher Scientific PA1-091) or rabbit anti-GFP isotype control antibody (Proteintech 50430-2-AP) for each co-IP reaction. Samples were then rotated overnight at 4°C to facilitate antibody-antigen binding. Protein G Dynabeads (Thermo Fisher Scientific 10004D) were prepared by rinsing 10.4 μl bead slurry (0.3125 mg beads) per reaction with IP lysis buffer (*Poitelon, et al., 2018*). The antibody-antigen complex was then captured by adding rinsed beads to each co-IP reaction and rotating for 1 hour at 4°C. Samples were then washed 3x five minutes at 4°C in IP lysis buffer, transferred to a fresh microcentrifuge tube during the final wash, and then eluted by boiling in 40 μl 2x Laemmli buffer for 10 minutes at 95°C for analysis by western blot (described in supplemental methods).

### Statistical analysis

All data collection and analyses were performed blind to the experimental conditions and animal genotype where applicable. No data were excluded from the analyses. Power analysis was not performed, but our sample sizes reflect those generally accepted. Figure legends report the statistical test used in each experiment. Graphs are presented as the mean average ± SEM. Statistical trends were reported as values where 0.05 < P < 0.1, while statistical significance was reported as values where P < 0.05. Data were analyzed using GraphPad Prism software v8.0.2.

### Reporting summary

The data supporting the findings reported in this study are available within this article and its associated supplementary information files. All raw data and biological resources can be made available from the corresponding author upon request.

## Acknowledgements

This work was funded by NIH NINDS R01 NS045630 (to M.L.F.), F30 NS118774 (to M.R.W.), R01 NS134493 (to S.B.) and R01 NS110627 (to Y.P.). We thank Gustavo Della-Flora Nunez and Emma R. Wilson for their technical assistance.

## Data Availability

The data supporting the findings of this study are available within the article and its supplementary information files. All data and biological resources (mouse lines, plasmids, etc.) are available from the corresponding authors upon reasonable request. Source data are provided with this paper.

## Author Contributions

M.R.W. and M.L.F. designed the research. M.R.W. carried out the majority of the experiments. D.S. quantified myelinated fibers and nerve cross-sectional area in semithin section images. S.P. and S.B. quantified P60 g-ratio in semithin section images. J.F. and Y.P quantify SC proliferation. M.L.F. quantified myelin abnormalities and amyelinated fibers in electron microscopy images. E.H. processed tissue for morphological analyses. M. Pellegatta, C.B., M. Palmisano, and S.F. performed the initial GST pulldown screen for Rac1 interactors. M.S. and F.E.P. performed mass spectroscopy. M.R.W., M.L.F., and Y.P. analyzed data. M.R.W. and Y.P. wrote the manuscript. F.S. critically reviewed the manuscript.

