## Supplemental Information for "The STRIPAK complex is required for radial sorting and laminin receptor expression in Schwann cells"

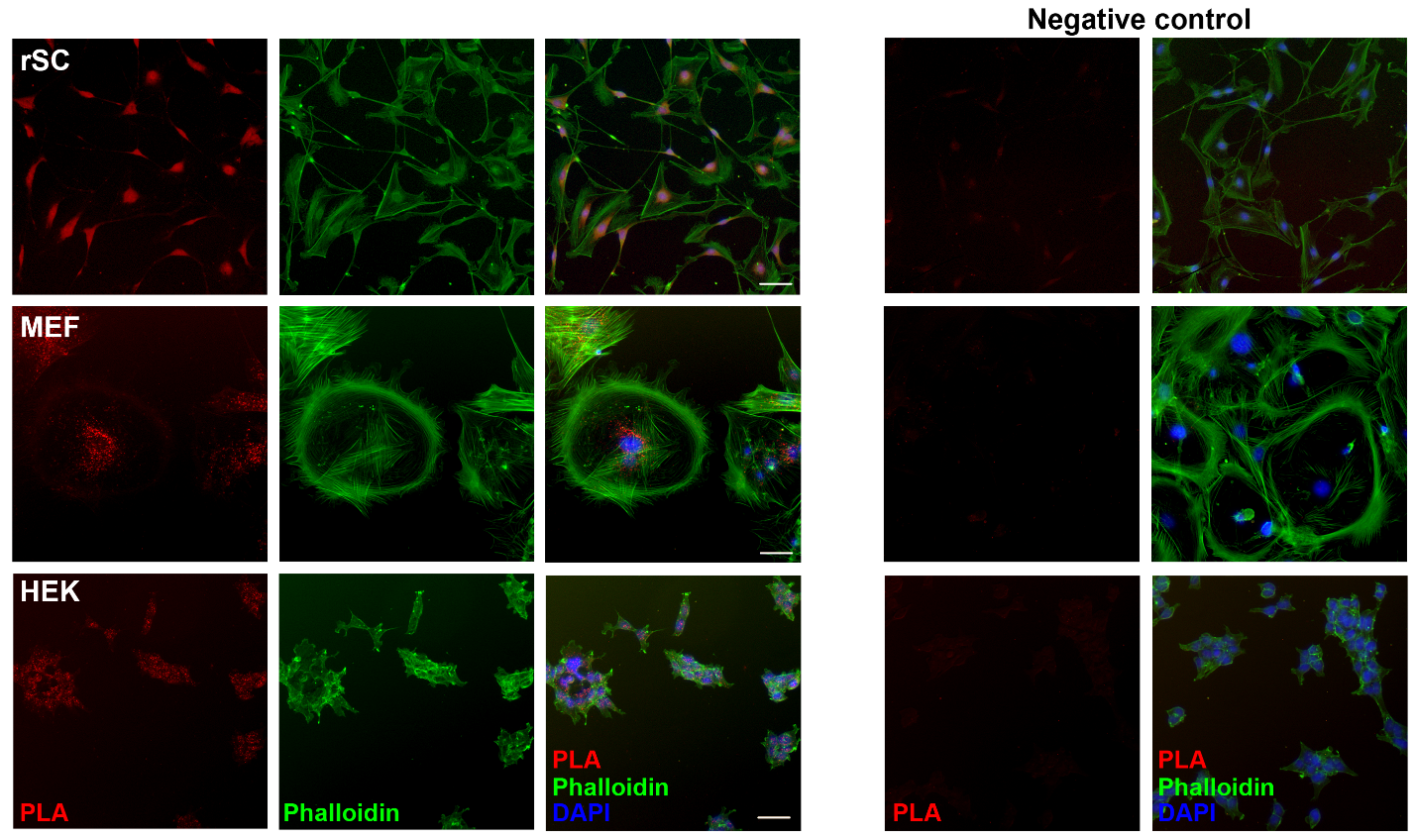

**Supplementary Figure 1: STRN3 and Rac1 are in close proximity in vitro.** Proximity ligation assay (PLA) technology permits the detection of protein-protein interactions in situ (< 40 nm). PLA analysis was performed in rat Schwann cells (rSC), mouse embryonic fibroblasts (MEF), and human embryonic kidney cells (HEK). The PLA analyses detected close localization (red dots) between STRN3 and Rac1 in the perinuclear region of rSC, MEF, and HEK. F-actin was stained with phalloidin (green) and nuclei were labeled with DAPI (blue). Negative controls lacking anti-STRN3 primary antibody did not produce PLA signals. Scale bar = 50 μm.

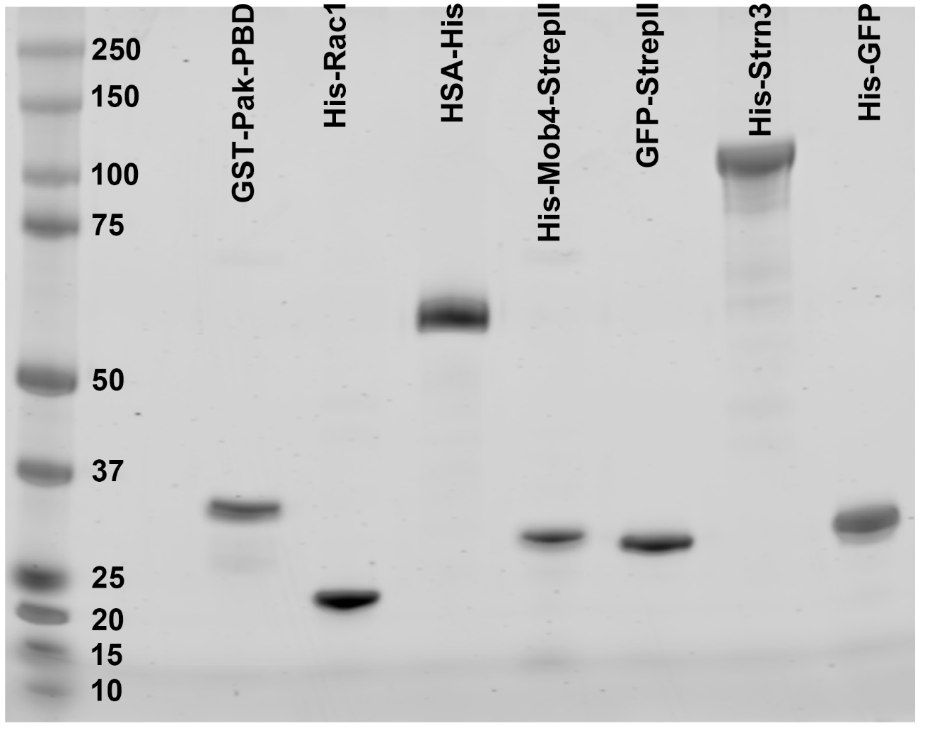

**Supplementary Figure 2: Protein purification produced high yield of recombinant proteins.** Gel electrophoresis for purified GST-PAK-PBD (Rac1/CDC42 binding domain (PDB) of p21 activated kinase 1 protein (PAK) labeled with a GST Tag; expected size 34kDa), His-Rac1 (Rac1 protein labeled with a His Tag, expected size 22 kDa), HSA-His (human serum albumin labeled with a His Tag; expected size 67 kDa), His-MOB4-StrepII (MOB4 protein labeled with a His Tag and a StrepII Tag, expected size 28 kDa), GFP-StrepII (green fluorescent protein labeled with a StrepII Tag, expected size 29 kDa), His-STRN3 (STRN3 protein labeled with a His Tag, expected size 95 kDa), His-GFP (green fluorescent protein labeled with a His Tag, expected size 33 kDa). Total proteins were stained with ReadyBlue.

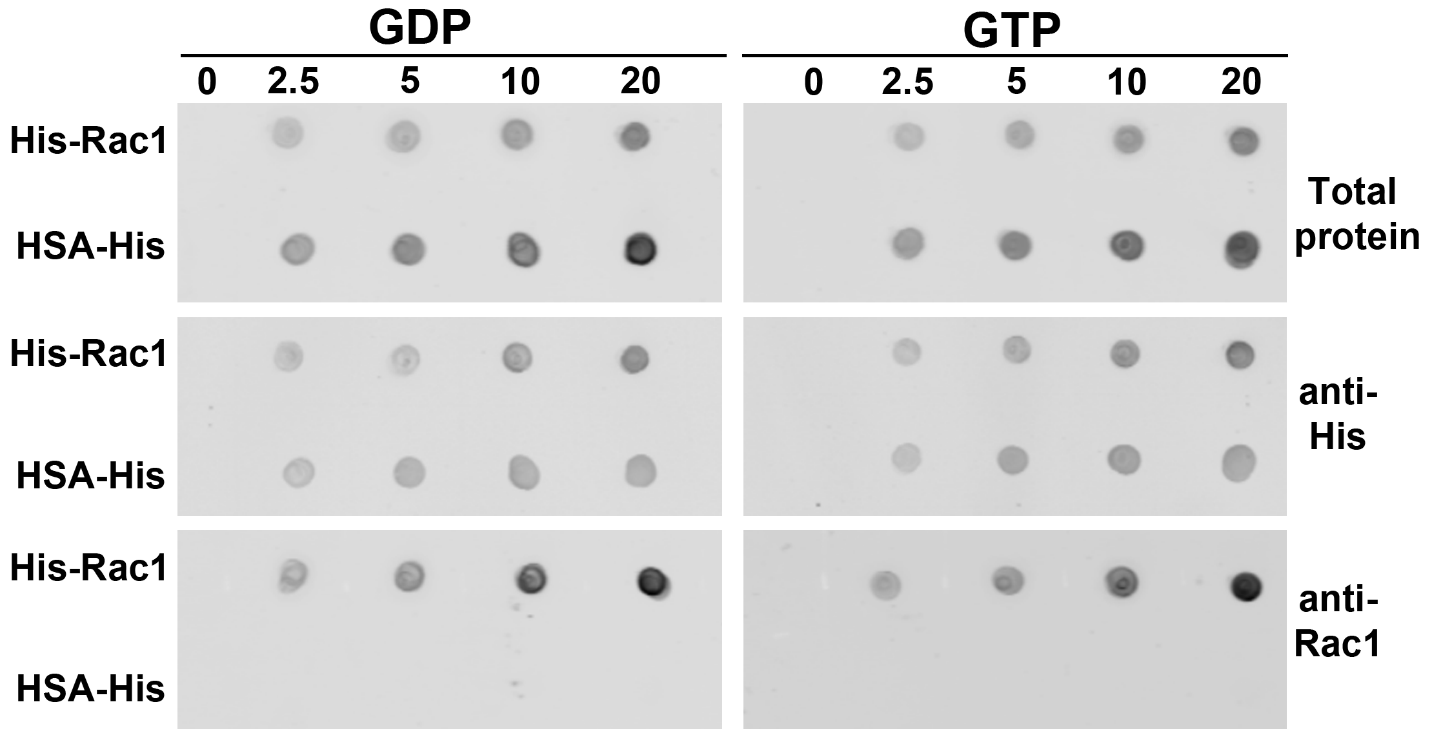

**Supplementary Figure 3: Representative dot blot membranes showing bait dots prior to incubation with prey proteins.** Bait proteins His-Rac1 (experimental) and HSA-His (negative control) were charged with either GDP or GTP and dotted in triplicate as a gradient (0, 2.5, 5, 10, and 20 pmoles). Membranes were separated by GDP/GTP charge state. Similar protein loading was observed for both GDP-charged and GTP-charged bait proteins by total protein staining and blotting with anti-His or anti-Rac1 antibodies. HSA-His has a higher molecular weight than His-Rac1, and therefore stains more intensely for total proteins when equal pmoles are loaded. Both bait proteins were labeled by anti-His blotting, but only His-Rac1 was labeled by anti-Rac1 blotting.

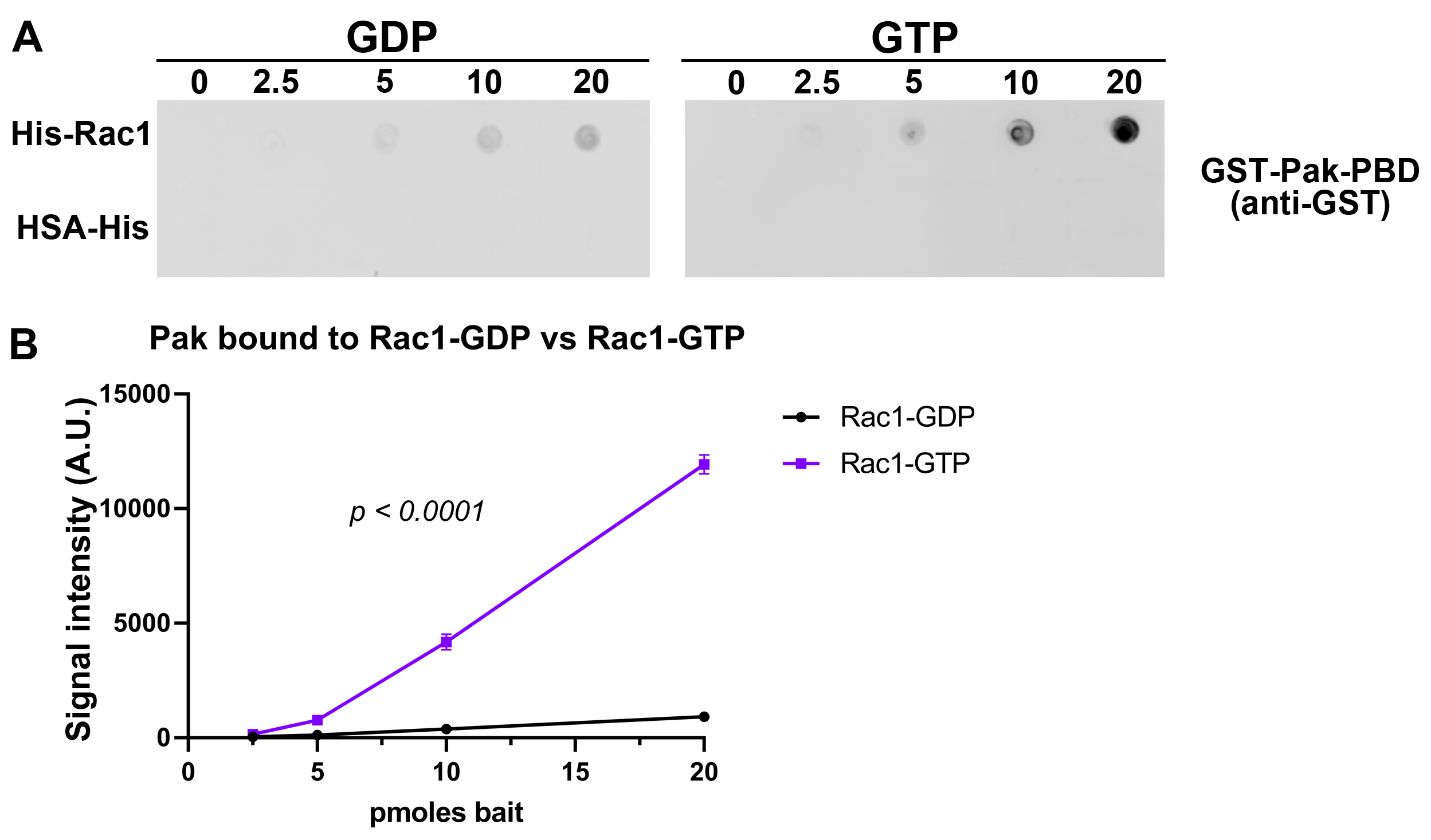

**Supplementary Figure 4: His-Rac1 maintains GDP/GTP charge state during dot blot procedure.** (**A**) Bait proteins His-Rac1 (experimental) and HSA-His (negative control) were charged with either GDP or GTP and dotted in triplicate as a gradient (0, 2.5, 5, 10, and 20 pmoles). Membranes were separated by GDP/GTP charge state. Dot blot membranes were incubated with prey protein GST-PAK-PBD and blotted for anti-GST. (**B**) Densitometry shows significantly more GST-PAK-PBD bound to His-Rac1-GTP as compared to His-Rac1-GDP. *N* = 3 individual dots per condition. Extra sum-of-squares F test. F (2, 20) = 421.1, *p* < 0.0001. Error bars indicate S.E.M. A.U.: Arbitrary Units.

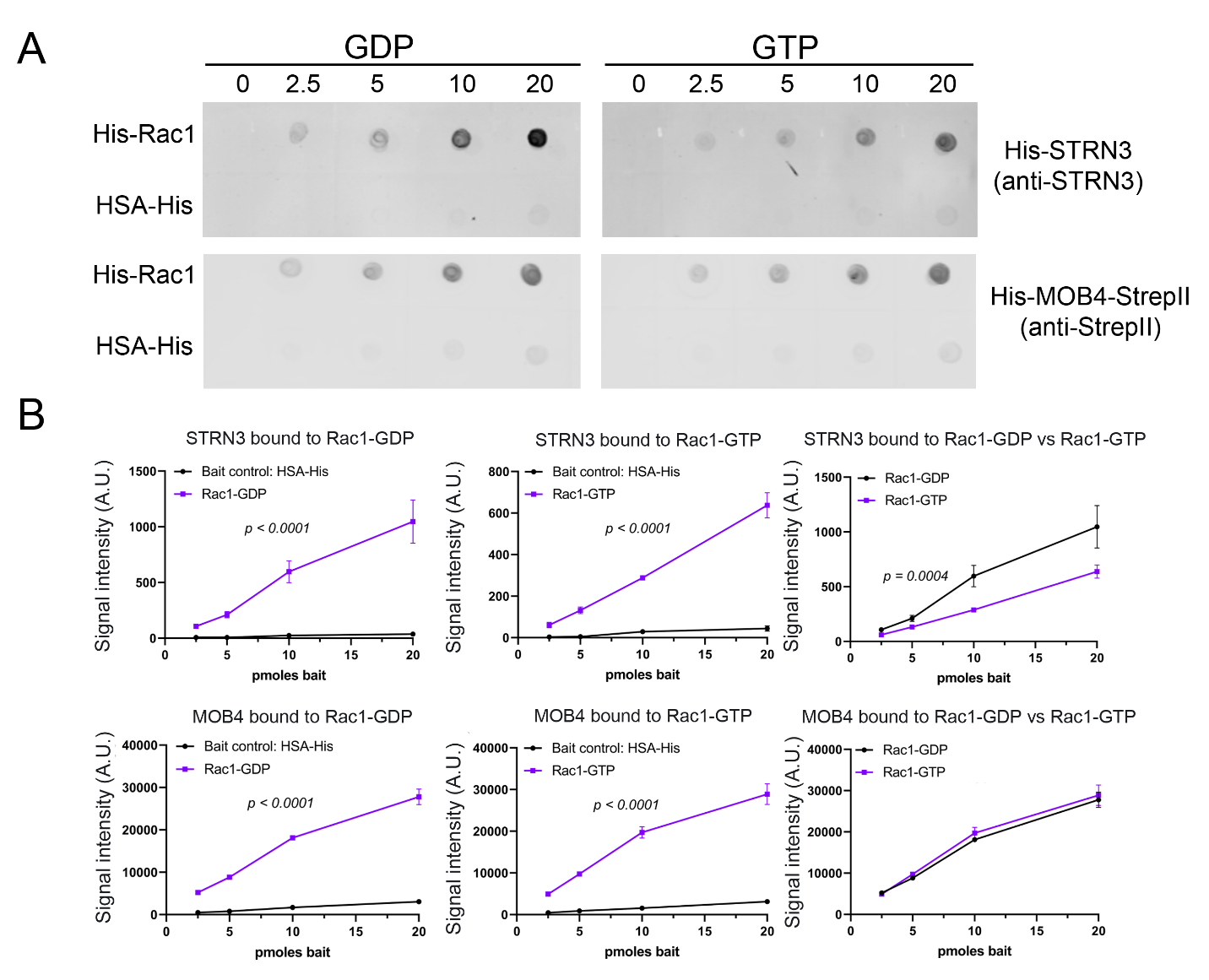

**Supplementary Figure 5: STRN3 interacts directly with Rac1.** (**A**) Bait proteins His-Rac1 (experimental) and HSA-His (negative control) were charged with either GDP or GTP and dotted in triplicate as a gradient (0, 2.5, 5, 10, and 20 pmoles). Membranes were separated by GDP/GTP charge state. Dot blot membranes were incubated with prey proteins His-STRN3 or His-MOB4-StrepII and blotted for anti-STRN3 or anti-StrepII, respectively. (**B**) Densitometry shows significantly more STRN3 bound to His-Rac1-GDP as compared to His-Rac1-GTP. In contrast, MOB4 bound to Rac1 independently of GDP/GTP charge state. *N* = 3 individual dots per condition. Extra sum-of-squares *F* test. A.U.: Arbitrary Units. Error bars indicate S.E.M. ** *p* < 0.01.

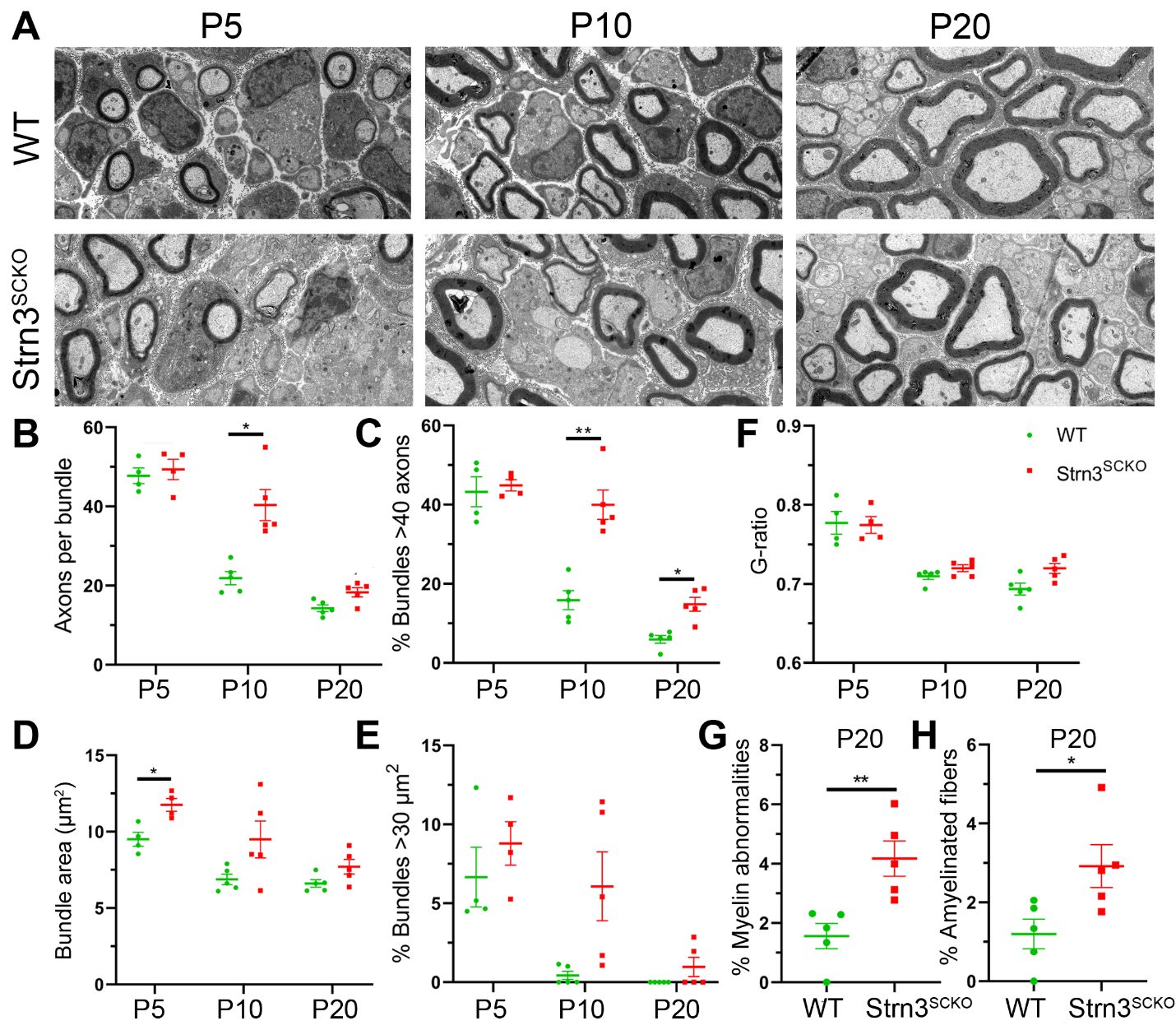

**Supplementary Figure 6: Ablation of *Strn3* in SCs results in mild radial sorting and myelin abnormalities.** (**A**) Representative electron microscopy images of WT and Strn3^SCKO^ sciatic nerves at P5, P10, and P20. Scale bar = 2µm. (**B**-**C**) Sciatic nerves of Strn3^SCKO^ mice have significantly more axons per unsorted, mixed-caliber axon bundle and significantly more bundles with many (>40) axons at P10. Two-way multiple comparisons ANOVA with Bonferroni post hoc test. Axons per bundle (*F* (1.743, 12.20) age = 137.3; *p* < 0.0001; *F* (1, 8) genotype = 11.11, *p* = 0.0103; *p*_P5_ > 0.9999, *p*_P10_ = 0.0190, *p*_P20_ = 0.0751). Bundles > 40 axons (*F* (1.299, 9.092) age = 95.32; *p* < 0.0001; *F* (1, 8) genotype = 24.09, *p* = 0.0012; *p*_P5_ > 0.9999, *p*_P10_ = 0.0031, *p*_P20_ = 0.0122). (**D**-**E**) Sciatic nerves of Strn3^SCKO^ mice have significantly larger axonal bundles at P5. Two-way multiple comparisons ANOVA with Bonferroni post hoc test. Bundle area (*F* (1.475, 10.33) age = 19.01; *p* = 0.0006; *F* (1, 8) genotype = 7.872, *p* = 0.0230; *p*_P5_ = 0.0327, *p*_P10_ = 0.2837, *p*_P20_ = 0.2592). Bundles > 30 µm^2^ (*F* (1.354, 14.89) age = 15.19; *p* = 0.0007; *F* (1, 22) genotype = 7.599, *p* = 0.0115; *p*_P5_ > 0.9999, *p*_P10_ = 0.1821, *p*_P20_ = 0.5626). (**F**) G-ratio (myelin thickness) is not significantly altered in sciatic nerves of Strn3^SCKO^ mice. Two-way multiple comparisons ANOVA with Bonferroni post hoc test. G-ratio (*F* (2, 11) age = 43.83; *p* < 0.0001; *F* (1, 11) genotype = 2.763, *p* = 0.1247; *p*_P5_ > 0.9999, *p*_P10_ > 0.9999, *p*_P20_ = 0.1170). (**G**) At P20, sciatic nerves of Strn3^SCKO^ mice have a significantly higher percentage of axons with myelin abnormalities, such as myelin infoldings, outfoldings, and decompaction. Unpaired two-tailed *t*-test (*t* = 3.568, df = 8, *p* = 0.0073). (**H**) At P20, sciatic nerves of Strn3^SCKO^ mice have a significantly higher percentage of sorted, amyelinated fibers. Unpaired two-tailed *t*-test (*t* = 2.605, df = 8, *p* = 0.0314). *N* = 4-5 samples per genotype. Error bars indicate S.E.M. (**B-F**) Two-way multiple comparisons ANOVA with Bonferroni post hoc test. **p* < 0.05, ** *p* < 0.01.

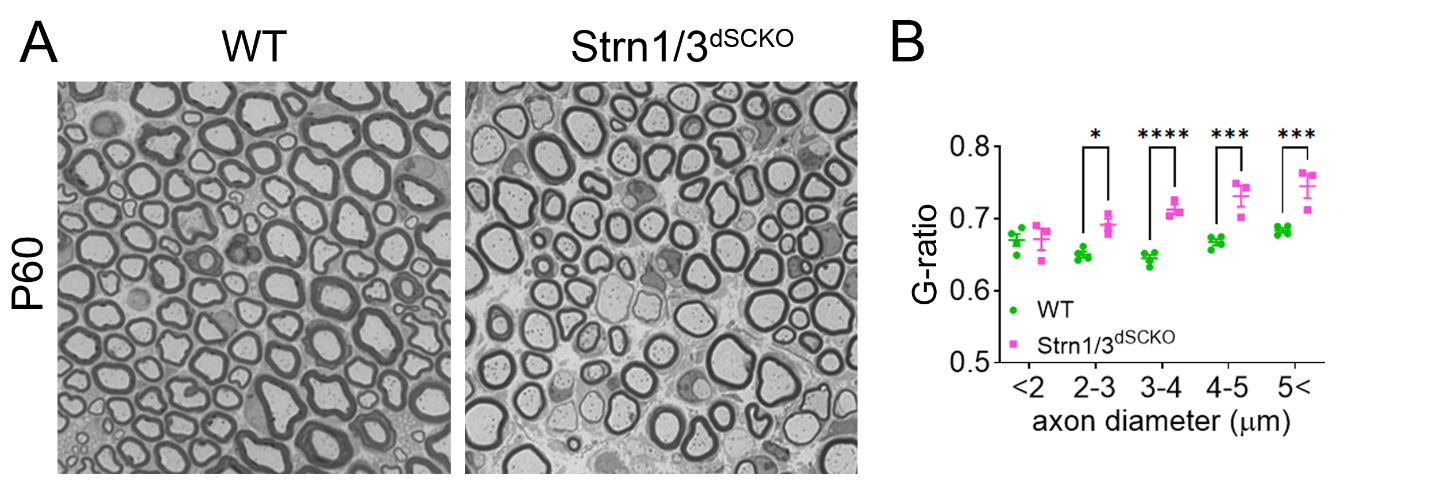

**Supplementary Figure 7: *Strn1* and *Strn3* ablation in SCs results in reduce myelin thickness.** (**A**) Representative semithin section images of WT and Strn1/3^dSCKO^ sciatic nerves at P60. Scale bar = 5µm. (**B**) G-ratio (myelin thickness) is significantly reduced in sciatic nerves of Strn1/3^dSCKO^ mice. Two-way multiple comparisons ANOVA with Bonferroni post hoc test. G-ratio (*F* (4, 25) axon diameter = 25.45; *p* < 0.0001; *F* (1, 25) genotype = 47.19, *p* < 0.0001; *p*_<2_ > 0.9999, *p*_2-3_ > 0.0144, *p*_3-4_ < 0.0001, *p*_4-5_ > 0.0001, *p*_5<_ < 0.0002. **p* < 0.05, *** *p* < 0.001, **** *p* < 0.0001.

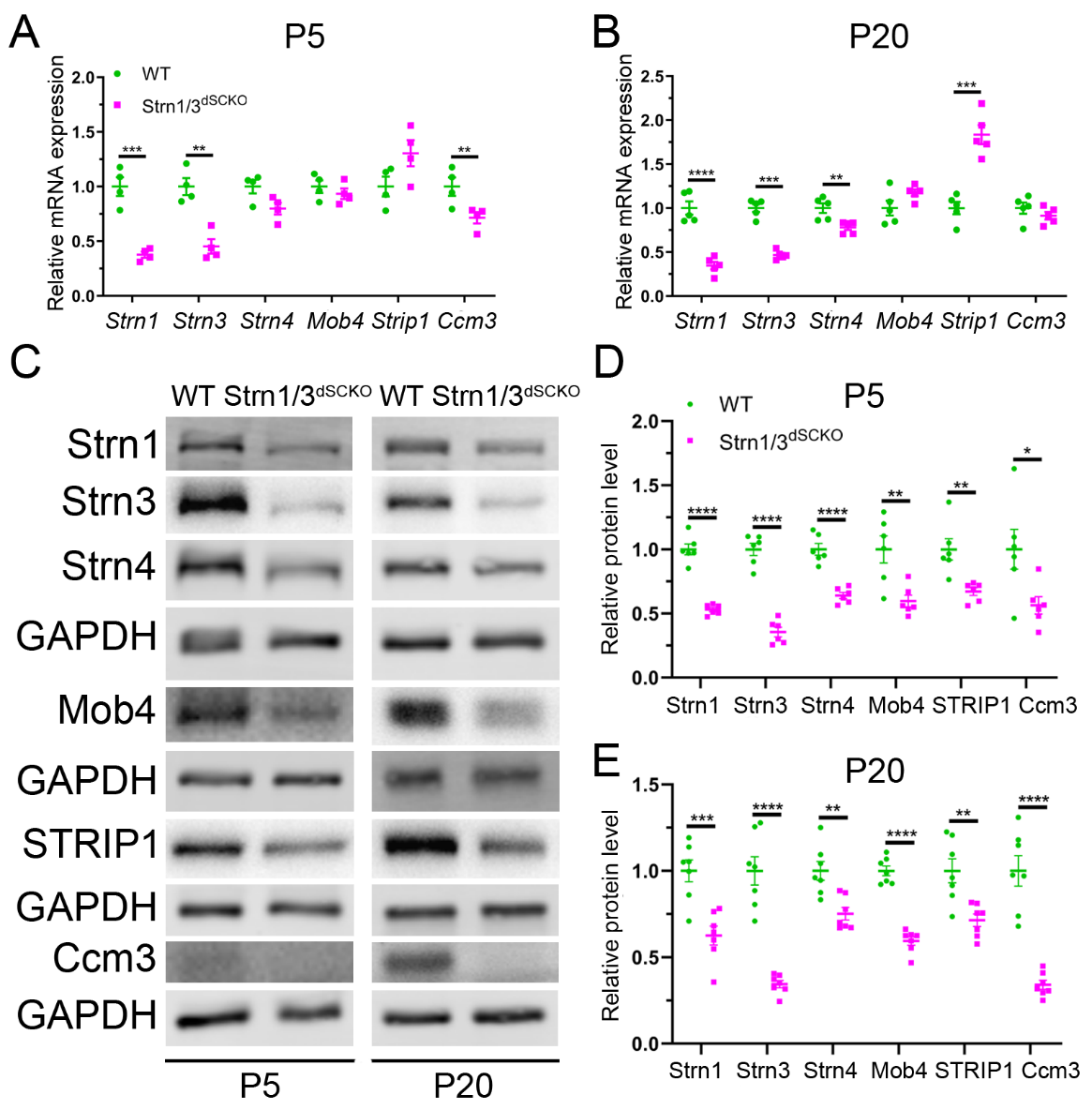

**Supplementary Figure 8: *Strn1* and *Strn3* ablation in SCs results in reduced levels of STRIPAK proteins.** (**A-B**) mRNA expression of several STRIPAK members from pooled sciatic nerves and brachial plexuses of WT and Strn1/3^dSCKO^ mice at P5 and P20. *Strn1* and *Strn3* mRNA levels are reduced at P5 and P20, *Ccm3* mRNA levels are reduced at P5, and *Strn4* mRNA levels are reduced at P20, and *Strip1* mRNA levels are increased at P20 in peripheral nerves of Strn1/3^dSCKO^ mice. *N* = 4-5 samples per genotype. P5 samples were pooled from 3-6 animals each. Unpaired two-tailed *t*-test. P5 [*Strn1* (*t* = 6.935, df = 6, *p* = 0.0004), *Strn3* (*t* = 5.385, df = 6, *p* = 0.0017), *Strn4* (*t* = 2.427, df = 6, *p* = 0.0514), *Mob4* (*t* = 0.8836, df = 6, *p* = 0.4109), *Strip1* (*t* = 2.032, df = 6, *p* = 0.0884), *Ccm3* (*t* = 2.906, df = 6, *p* = 0.0271)]. P20 [*Strn1* (*t* = 7.531, df = 8, *p* < 0.0001), *Strn3* (*t* = 10.48, df = 8, *p* < 0.0001), *Strn4* (*t* = 3.456, df = 8, *p* = 0.0086), *Mob4* (*t* = 0.1.858, df = 8, *p* = 0.1002), *Strip1* (*t* = 6.519, df = 8, *p* = 0.002), *Ccm3* (*t* = 1.140, df = 8, *p* = 0.2871)]. (**C**) Representative western blots of several STRIPAK proteins from pooled sciatic nerves and brachial plexuses of WT and Strn1/3^dSCKO^ mice at P5 and P20. (**D**-**E**) Densitometry analysis shows a reduction in STRN1, STRN3, STRN4, MOB4, STRIP1, and CCM3 protein levels in peripheral nerves of Strn1/3^dSCKO^ mice at P5 and P20. *N* = 6-7 samples per genotype. P5 samples were pooled from 3-6 animals each. Unpaired two-tailed *t*-test. P5 [STRN1 (*t* = 10.20, df = 10, *p* < 0.0001), STRN3 (*t* = 10.56, df = 10, *p* < 0.0001), STRN4 (*t* = 6.998, df = 10, *p* < 0.0001), MOB4 (*t* = 3.506, df = 10, *p* = 0.0057), STRIP1 (*t* = 3.718, df = 10, *p* = 0.0040), CCM3 (*t* = 2.585, df = 10, *p* = 0.0272)]. P20 [STRN1 (*t* = 4.497, df = 12, *p* = 0.0007), STRN3 (*t* = 7.770, df = 12, *p* < 0.0001), STRN4 (*t* = 3.842, df = 12, *p* = 0.0023), MOB4 (*t* = 10.84, df = 12, *p* < 0.0001), STRIP1 (*t* = 3.661, df = 12, *p* = 0.0033), CCM3 (*t* = 7.213, df = 12, *p* < 0.0001)]. Error bars indicate S.E.M. **p* < 0.05, ** *p* < 0.01, *** *p* < 0.001, **** *p* < 0.0001.

**Supplementary Methods**

**Genotyping.** Genomic DNA was used for genotyping as follows. Thermocycler conditions for all genotypes were 94°C for 5 minutes, (94°C for 15s, 65-56°C [-1°C per cycle] for 30s, 72°C for 40s) for 10 cycles, (94°C for 15s, 55°C for 30s, 72°C for 40s) for 30 cycles, 72°C for 5 minutes. PCR reaction mixtures were prepared with Quick Load Taq-Mix 2x (New England Biolabs M0271L). For MPZ-Cre, primers were A: 5’- CCACCACCTCTCCATTGCAC-3’, AS2: 5’- GCTGGCCCAAATGTTGCTGG-3’, 5MP2: 5’- TGTTGGCAACTTTGGATGTGT-3’, and P3P: 5’- TCAGCCAAGCCTTACCTTACT-3’, yielding bands of ~203 bp (internal control) and ~450 bp (transgene). For Strn1-floxed, primers were Strn1-F: 5’- TGAATTATTGGAGTTTTGTTTCAGACC-3’ and Strn1-ttR: 5’- GCACAGACAGACCTTCATGCTAACC-3’, yielding bands of ~531 bp (WT) and ~630 bp (floxed). For Strn3-floxed, primers were Strn3-F: 5’- ACCACAAAACAAGTGGTAGCTGAACC-3’ and Strn3-ttR: 5’- TGAAGGTGGTAGGAATACAGAATAGCC-3, yielding bands of ~684 bp (WT) and ~818bp (floxed). For Strn4 floxed, primers were Strn4-Fwd-3: 5’-CAGAGCAGCTGCTTGGCATAGAG-3’ and Strn4-Rev-3: 5’-CACAACCGTGCACACTGGTGC-3’, yielding bands of ~645bp (WT) and ~733bp (floxed). For Rac1-floxed, primers were Rac1-1: 5’- ATTTTGTGCCAAGGACAGTGACAAGCT-3’, Rac1-2: 5’- GAAGGAGAAGAAGCTGACTCCCATC-3’, and Rac1-3: 5’- CAGCCACAGGCAATGACAGATGTTC-3’, yielding bands of ~175bp (recombined), ~300bp (WT), and ~333bp (floxed).

**Dot blot.** For each charge reaction, bait proteins His-Rac1 (Cytoskeleton, Inc. RC01-XL) or His-human serum albumin (His-HSA, Abcam ab217817) negative control were diluted to 10 μM in 140 μl PD6 buffer (5 mM Tris-HCl pH 7.4, 12.5 mM NaCl_2_, 0.25 mM MgCl_2_, and 1:400 protease inhibitor cocktail) containing 1 mM GDP or GTPγS and incubated for 20 minutes at 37°C. Nitrocellulose membranes were labelled in pencil with a 1x1 cm grid pattern, and 2 μl dots of a serial dilution of His-Rac1-GDP/GTPγS or His-HSA-GDP/GTPγS charge reactions were dotted along the membranes. The serial dilution was as follows: blank charge reaction, 2.5 pmoles, 5 pmoles, 10 pmoles, and 20 pmoles. Rows along each membrane alternated between His-Rac1 and His-HSA dots, while membranes were separated by GDP vs GTPγS charge state. After laying protein dots, membranes were allowed to air dry for 15 minutes at room temperature. Membranes were rehydrated in TBS-T + 1 mM MgCl_2_, washed 3x with TBS-T + 1 mM MgCl_2_, and then blocked in 5% BSA in TBS-T + 1 mM MgCl_2_ for 1 hour at room temperature. Membranes were then incubated with prey proteins GST-PAK-PBD, His-Strn3 (MyBioSource MBS1348331), or His-Mob4-StrepII (prepared from Addgene expression plasmid # 31592, http://n2t.net/addgene:31592, RRID: Addgene_31592, a gift from Konrad Büssow) diluted to 100 nM in 5% BSA in TBS-T + 1 mM MgCl_2_ overnight at 4°C^[70]^. Membranes were then washed 3x with TBS-T and then immunolabelled and imaged according to a slightly modified version of the western blot protocol described in supplemental methods. Differences included reducing the primary antibody incubation time to 1 hour and performing the secondary antibody incubation at 4°C.

**Morphological assessment.** Mice were euthanized at the specified ages and then sciatic nerves were dissected, fixed in 2% glutaraldehyde, and stored at 4°C until processing. Nerves were then post-fixed in 1% osmium tetroxide, dehydrated by serial incubations of increasing ethanol concentration, and embedded in Epon resin with a transition solvent of propylene oxide. Embedded samples were then cut as semithin sections (1 μm) and stained with 2% toluidine blue for light microscopy or as ultrathin sections (80-85 nm) and stained with uranyl acetate and lead citrate for electron microscopy. For analysis of g-ratio, myelinated fiber density, total myelinated fibers per nerve, and nerve cross-sectional area, images were acquired with the 100x objective of a Leica DM6000 microscope and Q-Capture Pro V7.0.4324.5 software (QImaging, Inc). G-ratio was subsequently calculated using semi-automated Leica QWin software (Leica Microsystem). For myelinated fiber density, total myelinated fibers per nerve, and nerve cross-sectional area, the 100x images were stitched together with PTGui software v.10 (New House Internet Services BV) to assemble a full nerve cross-section. Morphological measurements were made using Fiji software v1.52h (ImageJ). Transmission electron micrographs at 2900x magnification were acquired and then analyzed by Fiji software to quantify unsorted axon bundle size, axons per unsorted bundle, % amyelinated axons, and % axons with myelin abnormalities (infoldings, outfoldings, and decompaction).

**Cell culture.** Primary rat SCs (rSC) were isolated from P3 rat sciatic nerves. Nerves were dissected, dissociated with 130 U/ml type I collagenase (Fischer Scientific NC9633623), and cells were then plated on poly-L-lysine (PLL, Sigma Aldrich P-5899)-coated tissue culture dishes. Plates were coated with PLL by incubation with 0.01 mg/ml PLL in water for 30 minutes at room temperature followed by washing with phosphate buffered saline (PBS). Contaminant fibroblasts were killed to achieve 99-100% rSC culture purity by treatment with cytosine β-D-arabinofuranoside hydrochloride (Sigma-Alrich C6645) followed by two treatments of mouse anti-Thy1.1 monoclonal antibody (Bio-Rad MCA045G) and rabbit complement. Cultures were limited to five passages in media containing high glucose DMEM (Thermo Fisher Scientific 11965-118) supplemented with 10% fetal bovine serum (FBS, Thermo Fisher Scientific 10437028), 2 mM L-glutamine (Thermo Fisher Scientific 25030-081), 100 U/mL penicillin + 100 μg/ml streptomycin (Thermo Fisher Scientific 15140-122), 2 ng/ml NRG1 (R&D Systems 396-HB-050), and 2 μM forskolin (EMD Millipore 344270). Primary mouse SCs (mSC) were prepared from pooled sciatic nerves of adult mice aged P45-60. Nerves were dissected, placed in ice-cold Leibovitz’s L-15 media (Thermo Fisher Scientific 11415064) supplemented with 100 U/mL penicillin + 100 μg/ml streptomycin, and stripped of epineurium and other contaminant tissues. Cleaned nerve explants were then transferred into tissue culture dishes (6-8 nerves / 35 mm dish) containing nerve explant / mSC media (identical to rSC media except for NRG1 increased to 10 ng/ml). Nerve explants were incubated at 37°C, 5% CO_2_ for 7 days with media changes every other day to allow for formation of repair SCs. Cells were then enzymatically dissociated by placing explants in L-15 media containing 2.5 mg/ml dispase II (Sigma-Aldrich D4693), 130 U/ml type I collagenase, and 5 mM HEPES while incubating at 37°C for three hours with gentle rocking at 40 rpm. Tissue was then mechanically dissociated using fire-polished glass pipettes followed by passage through a 70 μM cell strainer. The cell suspension was then centrifuged, the resulting pellet was resuspended in mSC media, and cells were plated on tissue culture plates coated with PLL and laminin 111 (Krackler/Sigma-Aldrich L2020). Plates were coated with PLL+laminin by incubation with 0.01 mg/ml PLL in water for 30 minutes at room temperature, washing with PBS, incubation with 0.01 mg/ml laminin 111 in PBS for 1 hour at room temperature, incubation with mSC media for 30 minutes at room temperature, and then washing with PBS. Resultant mSC cultures were passaged once as appropriate for each experiment. Mouse embryonic fibroblasts (MEFs) were isolated from E12.5-14.5 mouse embryos. Embryos were dissected from pregnant female mice and placed in a tissue culture dish with ice cold PBS. Yolk sac, head, heart, and liver were removed. Embryos were then placed into a fresh tissue culture dish with 0.25% trypsin-EDTA (Thermo Fisher Scientific 15090046) in PBS, minced, incubated at 37°C for 10 minutes, and then pipetted several times to obtain a near single-cell suspension. Cells were transferred to a dish coated with collagen I (Corning 354236) and allowed to adhere by incubating overnight at 37°C, 5% CO_2_ in MEF/HEK media (high glucose DMEM supplemented with 10% FBS and 100 U/mL penicillin + 100 μg/ml streptomycin). Collagen coating was performed by incubating dishes with 0.01 mg/ml collagen I in PBS for 1 hour at room temperature. After adhering overnight, residual debris was washed away wish PBS. MEFs were limited to 10 passages.

**Proximity ligation assay.** Cells were plated on 96-well plates at 8000 cells / well and allowed to adhere overnight at 37°C, 5% CO_2_. rSCs were plated on wells coated with PLL + laminin 111 (as described above), while MEF and human embryonic kidney (HEK) cells were plated on PLL + collagen I. The procedure for PLL + collagen I coating was identical to that for PLL + laminin 111 coating, except that laminin 111 was substituted for collagen I diluted to 0.01 mg/ml in PBS. rSCs were incubated in rSC media while MEFs and HEKs were incubated in MEF/HEK media (described above). Cells were fixed by incubating with 4% PFA in PBS at room temperature for 20 minutes. Cells were then permeabilized and stained for F-actin by incubating with phalloidin-488 1:200 (Cytoskeleton, Inc. PHDG1) in PBS and 0.5% Triton X-100 for one hour at room temperature. Samples were then blocked and nuclei were stained by incubating with DAPI in Duolink blocking solution (Duolink In Situ Detection Reagents Red kit, Sigma-Aldrich DUO92008) for one hour at room temperature. Samples were then immunolabeled by incubating with primary antibodies in Duolink antibody diluent overnight at 4°C, washing 3x in Duolink wash buffer A, incubating with Duolink secondary antibody probes (anti-rabbit Plus and anti-mouse Minus) each diluted 1:5 in Duolink antibody diluent overnight at 37°C, and then washing 3x in Duolink wash buffer A. Duolink probes were ligated by incubating with Duolink ligase enzyme diluted 1:40 in Duolink ligase buffer for 1 hour at 37°C and then washing 3x in Duolink wash buffer A. Ligated DNA probes were amplified by rolling circle amplification (RCA) by incubation with Duolink polymerase diluted 1:80 in Duolink amplification buffer overnight at 37°C. Samples were then washed 3x with Duolink wash buffer B, 1x with 1:100 Duolink wash buffer B, 1x with PBS, and then imaged in PBS. The following primary antibodies were used: mouse anti-striatin-3 (Novus Biologicals NB110-74572) and rabbit anti-Rac1 (Proteintech 24072-1-AP). Immunofluorescent images were acquired as described above.

**RNA extraction and RT-qPCR analysis.** Sciatic nerves and brachial plexuses were dissected at the indicated ages, contaminant tissue was removed, samples were snap frozen in liquid nitrogen, and then stored at -80°C. Nerves were then pulverized, total RNA was isolated using Trizol (Thermo Fisher Scientific 15596026), and cDNA was prepared using the Superscript III kit (Thermo Fisher Scientific 18080051). For each reverse transcription reaction, 350 μg RNA and 5 μM of oligo(dT) were used. For SYBR qPCR reactions, SYBR green qPCR Master Mix (Thermo Fisher Scientific 4309155) was used for the following primer sets. For Itga6, primers were Fwd: 5’- cctgaaagaaaataccagactctca-3’, Rev: 5’-ggaacgaagaacgagagagg-3’. For Itgb1, primers were Fwd: 5’-CAACCACAACAGCTGCTTCTAA-3’, Rev: 5’-TCAGC CCTCTTGAATTTTAATGT-3’. For Itgb4, primers were Fwd: 5’-CTTGGTCGCCGTCTGGTA-3’, Rev: 5’- TCGAAGGACACTACCCCACT-3’. For Dag1, primers were Fwd: 5’-ctgctgctgctccctttc-3’, Rev: 5’-gcagtgttgaaaaccttatcttcc-3’. For GAPDH, primers were Fwd: 5’-CAACTCCCTCAAGATTGTCAGCAA-3’, Rev: 5’-GGCATGGACTGTGGTCATGA-3’. For Taqman qPCR reactions, Taqman Universal PCR Master Mix (Thermo Fisher Scientific 4364338) was used for the following Thermo Fisher Scientific Taqman assay gene probes: Ccm3 (Mm00727342_s1), Egr2/Krox20 (Mm00456650_m1), GAPDH (Mm99999915_g1), Mst1 (Mm00451755_m1), Mst2 (Mm00490480_m1), Mob4 (Mm00481145_m1), Oct6/Pou3f1 (Mm00843534_s1), Strip1 (Mm00463714_m1), Strn1 (Mm00448910_m1), Strn3 (Mm00453492_g1), Strn4 (Mm00467125_m1), Taz (Mm01289583_m1), and Yap (Mm01143263_m1). SYBR and Taqman qPCR assays were normalized to GAPDH reference gene. All qPCR assays were performed using a Bio-Rad CFX96/384 RT-qPCR machine with the following thermocycler protocol: 95°C for 10 minutes following by 40 cycles of (95°C for 15 seconds and 60°C for 1 minute). Data were analyzed using threshold cycle (Ct) and 2(−ΔΔCt) with the average expression of WT control animals normalized to 1.

**Western blot.** Sciatic nerves and brachial plexuses were dissected at the indicated ages, contaminant tissue was removed, samples were snap frozen in liquid nitrogen, and then stored at -80°C. Nerves were then pulverized and lysed in RIPA lysis buffer (50 mM Tris pH 7.4, 150mM, NaCl, 1% IGEPAL CA-630, 0.1% SDS, 0.5% sodium deoxycholate, 5mM EDTA, 1mM EGTA, 1 mM NaF, 1:100 protease inhibitor cocktail (Sigma-Aldrich P8340), 1:100 phosphatase inhibitor cocktail 2 (Sigma-Aldrich P5726), and 1:100 phosphatase inhibitor cocktail 3 (Sigma-Aldrich P0044)). Cells were lysed directly from plates using RIPA lysis buffer. Following lysis, samples were sonicated for two cycles of 30 seconds at 70% power and centrifuged at 13,200 rcf for 15 minutes at 4°C. Protein concentration of the supernatant was quantified by BCA assay (Thermo Fisher Scientific 23225). Loading samples for each experiment were prepared at equal concentrations by diluting supernatant with RIPA lysis buffer and 4x Laemmli sample buffer (Bio-Rad 1610747, supplemented with β-mercaptoethanol as per manufacturer’s instructions). Loading samples were electrophoresed through an SDS-polyacrylamide hydrogel and then transferred onto a PVDF membrane. Membranes were then blocked with 5% bovine serum albumin (BSA, Sigma-Aldrich A4161) in TBS-T (1x tris-buffered saline + 0.1% Tween-20) and immunolabelled with primary antibody in 5% BSA in TBS-T overnight at 4°C with gentle rocking. Next, membranes were washed with TBS-T, incubated with secondary antibody in 5% BSA in TBS-T for one hour at room temperature, and washed again with TBS-T. The following primary antibodies were used: mouse anti-β-Actin 1:1000 (Santa Cruz sc-47778), mouse anti-β-tubulin 1:2000 (Sigma-Aldrich T4026), mouse anti-β-dystroglycan 1:500 (Leica Biosystems NCL-b-DG), rabbit anti-Ccm3 1:250 (Proteintech 10294-2-AP), rabbit anti-Cdc42 1:250 (Cell Signaling 2462), rabbit anti-Egr2/Krox20 1:500 (kindly shared by D. Meijer), rabbit anti-GAPDH 1:10,000 (Sigma Aldrich G9545), rabbit anti-GST tag 1:1000 (Proteintech 80006-1-RR), mouse anti-GST tag 1:1000 (Santa Cruz sc-138), rabbit anti-His tag 1:1000 (Cell Signaling 2365S), rabbit anti-His tag 1:50,000 (Cell Signaling 2366T), goat anti-integrin α6 1:250 (Santa Cruz sc-6597), rabbit anti-integrin β1 1:250 (Cell Signaling 4706), rat anti-integrin β4 1:250 (Abcam ab25254), mouse anti-Mob4 1:250 (Santa Cruz sc-137229), rabbit anti-Mst1 1:500 (Cell Signaling 3682), rabbit anti-Mst2 1:500 (Abcam ab52641), rabbit anti-p-Mst1/2 1:250 (Proteintech 28953-1-AP), rabbit anti-NF2 1:500 (Cell Signaling 6995), rabbit anti-p-NF2 1:500 (Cell Signaling D5A4I), rabbit anti-Oct6 1:500 (kindly shared by D. Meijer), rabbit anti-Pak1 1:500 (Cell Signaling 2602), rabbit anti-p-Pak1 1:500 (Cell Signaling 2601), mouse anti-Rac1 1:250 (EMD Millipore 05-389), mouse anti-Rac1 1:250 (Cytoskeleton, Inc. ARC03), rabbit anti-Rac1 1:500 (Thermo Fisher Scientific PA1-091), rabbit anti-Rac1 1:500 (Proteintech 24072-1-AP), rabbit anti-Sox10 1:250 (Cell Signaling 89356), mouse anti-StrepII tag 1:250 (Thermo Fisher Scientific MA5-37747), mouse anti-STRIP1 1:250 (Origene TA502314), mouse anti-striatin-1 (Santa Cruz sc-136084), rabbit anti-striatin-3 1:100 (Atlas Antibodies HPA004636), mouse anti-striatin-3 (Novus Biologicals NB110-74572), rabbit anti-striatin-4 (GeneTex GTX133282), rabbit anti-Yap 1:250 (Cell Signaling 4912), rabbit anti-Yap 1:250 (Cell Signaling 14074), rabbit anti-Yap/Taz 1:250 (Cell Signaling 8418), rabbit anti-p-Yap 1:250 (Cell Signaling 13008), rabbit anti-p-Taz 1:250 (Cell Signaling 59971). The following secondary antibodies were used: donkey anti-rabbit 680 1:20,000 (Li-Cor 926-68073), goat anti-rabbit 800 1:20,000 (Li-Cor 926-32211), goat anti-mouse IgG1 680 (Li-Cor 926-68050), goat anti-mouse IgG2b 800 1:20,000 (Li-Cor 926-32352), goat anti-rat 680 1:20,000 (Li-Cor 926-68029), goat anti-rat 800 1:20,000 (Li-Cor 926-32219), donkey anti-goat 680 1:20,000 (Li-Cor 926-68024), donkey anti-goat 800 1:20,000 (Li-Cor 926-32214), donkey anti-rabbit HRP 1:20,000 (Jackson ImmunoResearch 711-035-152), goat anti-mouse IgG1 HRP 1:20,000 (Thermo Fisher Scientific A10551), goat anti-mouse IgG2a HRP 1:20,000 (SouthernBiotech 1080-05), goat anti-mouse IgG2b HRP 1:20,000 (Thermo Fisher Scientific M32407), goat anti-mouse Fc HRP 1:20,000 (Sigma-Aldrich A2554), goat anti-mouse kappa light chain HRP 1:20,000 (Jackson ImmunoResearch 115-035-174), goat anti-rat light chain HRP 1:20,000 (Jackson ImmunoResearch 112-035-175), donkey anti-goat HRP 1:20,000 (Jackson ImmunoResearch 705-036-147). Membranes were either imaged directly with Li-Cor Odyssey CLx infrared imaging system or imaged using a ChemiDoc XRS system after developing with ECL Select (GE Healthcare). Bands were quantified with either Image Studio Lite 5.2 software (Odyssey) for infrared blots or Image Lab 6.0 software (Bio-Rad) for chemiluminescent blots. GAPDH was used as a loading control.

**Immunofluorescence.** Samples were first fixed by incubation with 4% paraformaldehyde (PFA) in PBS on ice for 30 minutes (nerves) or 20 minutes (cells). Nerve samples were then cryprotected by incubating in 20% sucrose in PBS overnight at 4°C, embedded in tissue freezing medium, and cut into 8 μM thick sections onto glass slides. Samples were blocked and permeabilized by incubation with blocking buffer containing 20% FBS, 1% BSA, and 1% Triton X-100 (Krackler/Sigma-Aldrich T8787) in PBS for one hour at room temperature. Immunolabeling of samples was accomplished by incubation with primary antibodies in blocking buffer overnight at 4°C, incubation with secondary antibodies for one hour at room temperature, counterstaining with DAPI (Krackler/Sigma-Aldrich 45-D9542-5MG) for 10 minutes at room temperature, and mounting slides or coverslips with Vectashield (Vector Laboratories H-1000-10). Where indicated, rhodamine-conjugated phalloidin 1:200 (Cytoskeleton, Inc. PHDR1) was incubated with secondary antibodies to stain for F-actin. The following primary antibodies were used: rabbit anti-striatin-3 1:100 (Atlas Antibodies HPA004636), mouse anti-striatin-3 (Novus Biologicals NB110-74572), rabbit anti-Sox10 1:100 (Cell Signaling 89356), chicken anti-neurofascin 1:1000 (R&D Systems AF3235), and rat anti-integrin α6 1:100 (Thermo Fisher Scientific 14-0495-85). The following secondary antibodies were used: donkey anti-rabbit 488 1:1000 (Jackson ImmunoResearch 711-545-152), donkey anti-rabbit Cy3 1:500 (Jackson ImmunoResearch 711-166-152), goat anti-rabbit 594 1:500 (Invitrogen A32740), goat anti-mouse IgG1 488 1:1000 (Thermo Fisher Scientific A21121), goat anti-mouse IgG1 Cy3 1:500 (Jackson ImmunoResearch 115-165-205), donkey anti-chicken 488 1:1000 (Jackson ImmunoResearch 703-545-155), donkey anti-rat Cy3 1:500 (Jackson ImmunoResearch 712-165-153). Immunofluorescence images were acquired using a Leica SP5II confocal microscope running LAS AF 2.7.9723.3 software.
